# FTO regulates centrosome function and mitotic fidelity in cancers with chromosome instability (CIN)

**DOI:** 10.64898/2026.03.22.713449

**Authors:** Lea Cohen-Attali, Yelena Mostinski, Guy Biber, Daniela Kaludjerski, Mor Hanan, Reut Eshel, Alexandra Polyansky, Igor Pery, Aric Orbach, Rotem Karni, Amir Mor

## Abstract

Centrosomes are essential for spindle assembly and accurate chromosome segregation during mitosis. We found that the RNA demethylase FTO localizes to centrosomes in both human and mouse cells. Disruption of FTO function through a newly developed small-molecule inhibitor impaired spindle organization and centrosome function. This was evidenced by chromosome misalignment during mitosis, prometaphase arrest, followed by mitotic slippage or apoptosis. Mechanistically, we found that FTO is essential for intact localization of NuMA and KIFC1 proteins required for chromosome localization during mitosis. Importantly, cancers characterized by chromosomal instability (CIN) showed increased sensitivity to FTO inhibition compared to normal cells. These findings underscore FTO as a promising therapeutic target in CIN cancer subsets.

## Background

The centrosome, composed of a pair of centrioles surrounded by pericentriolar material, is the primary microtubule-organizing center of animal cells and plays a central role in spindle assembly during mitosis^1,2^. Synchronously with DNA replication, centrosomes duplicate once, during the S phase of the cell cycle. Upon entry into mitosis, the duplicated centrosomes migrate to opposite poles of the cell and nucleate the bipolar spindle, ensuring accurate microtubule-kinetochore attachment and faithful chromosome segregation^3^.

N6-methyladenosine (m6A) is the most abundant internal modification in eukaryotic RNA and has emerged as a key regulator of RNA metabolism. This modification is deposited by “writers” (the METTL3 (Methyltransferase-like 3)-METTL14 (Methyltransferase-like 14)-WTAP (Wilms’ tumor 1-associating protein) methyltransferase complex), removed by “erasers” (the demethylases FTO (Fat Mass and Obesity-associated protein) and ALKBH5 (AlkB Homolog 5, RNA Demethylase)), and interpreted by “readers” (including YTH-domain proteins, splicing factors and the translation machinery) that link m6A marks to downstream RNA processes^4,5^. Emerging evidence indicates that dynamic m6A modification plays a critical role in regulating mitotic entry, spindle assembly, and chromosome segregation^6,7,8^. However, the specific contribution of the FTO RNA demethylase to these processes remains poorly understood. While FTO has been shown to be essential in mouse spermatogonia, where its knockout leads to chromosome instability and G2/M arrest^9^, its relevance to mitotic regulation in human cells, both in normal physiology and disease, has yet to be fully elucidated.

FTO is an AlkB family dioxygenase that catalyzes the demethylation of N6-methyladenosine (m6A) and N6,2’-O-dimethyladenosine (m6Am) in RNA transcripts^10^. The protein contains a characteristic double-stranded β-helix fold that binds Fe(II) and α-ketoglutarate as cofactors for its catalytic activity^11^. FTO was documented to localize primarily to the nucleus, with a smaller fraction present in the cytoplasm^12^. Beyond its initial association with energy metabolism, FTO has emerged as an important regulator in cancer biology^13^. Its overexpression has been documented in multiple malignancies, including acute myeloid leukemia (AML) and various hematologic and solid tumors^14^. However, its specific role in cancers characterized by chromosomal instability remains largely unexplored.

Chromosomal instability (CIN) is a hallmark of cancer characterized by ongoing errors in chromosome segregation, leading to aneuploidy and genetic heterogeneity within tumors^15,16^. A particularly significant manifestation of CIN is whole genome doubling (WGD), an event in which cells double their DNA content without completing cytokinesis, leading to tetraploidy^17,18^. Unattached or improperly attached kinetochores can delay mitotic exit and, depending on the cell type, lead to apoptosis or mitotic slippage, an abnormal process that can trigger WGD^18,19^. Importantly, cancers exhibiting CIN or WGD display heightened sensitivity to perturbations of factors involved in chromosome alignment during mitosis, making these molecular players attractive therapeutic targets in these cancers^20–24^.

Here we report on newly developed small-molecule FTO inhibitors that induce misalignment of chromosomes, leading to prometaphase arrest and subsequent mitotic slippage or apoptosis. This phenotype was validated by siRNA-mediated FTO knockdown, which predominantly resulted in mitotic slippage. Mechanistically, we demonstrated that FTO is an integral component of the centrosome. Rescue studies with our inhibitor further revealed that FTO inhibition causes mislocalization of key centrosomal proteins essential for chromosome fidelity, including NuMA and KIFC1^25^. Consistent with previous findings for KIFC1 and other Kinesins^26^, cancers exhibiting CIN were highly sensitive to FTO inhibition, whereas normal cells were significantly less affected. These results highlight FTO inhibition as a selective therapeutic strategy for cancers with CIN.

## Results

### Discovery and development of novel FTO inhibitors

FTO has been identified as a key target in several cancers^14,27,28^; however, existing FTO inhibitors have limited specificity^29^, inadequate potency, and poor pharmacokinetic properties, which have hindered their advancement beyond early experimental stages^13^. To overcome these limitations, we initiated a drug discovery program to develop a potent FTO inhibitor for cancer treatment (Figure 1A). From an internal virtual screening campaign, followed by structure-activity relationship studies (SAR), 780 hits were assessed for their ability to inhibit recombinant FTO (for full details, see Methods section and Supplementary Figure S1A,B). 41 validated hits, inhibiting recombinant FTO at IC_50_<1µM, were further tested for their ability to bind cellular human FTO as measured by cellular thermal shift assay (CETSA) (For full details, see Methods section and Supplementary Figure S1C,D). Additionally, we tested their ability to inhibit MV-411 AML cancer cells that were previously shown to be sensitive to FTO inhibition^28^. PBMCs from a healthy donor were chosen to estimate the unspecific cytotoxicity of hit compounds. A promising series of inhibitors was obtained and optimized through further SAR studies, culminating in the identification of potential lead candidates, RNB-557 and RNB-637 (Figure 1B–E). Notably, SAR studies yielded inactive control isomer molecules, RNB-558 and RNB-638 (Figure 1B,D), that used as efficient tools for on-target effect validation. The active compounds RNB-557 and RNB-637 effectively inhibited AML cancer cell growth, whereas the inactive isomer molecules, RNB-558 and RNB-638 showed no inhibitory effect in both cell-free recombinant-FTO assay and in cells (Figure 1E,G and Supplementary Figure S1E). In addition, RNB-557 and RNB-637 bind mouse FTO, as demonstrated by CETSA performed in the B16 melanoma cell line, confirming cross-species FTO reactivity (Supplementary Figure S1F). Furthermore, molecules obtained through SAR analysis of the RNB-557/637 series exhibited a strong correlation between CETSA binding scores, recombinant FTO inhibition IC_50_ values (Figure 1F) and MV-411 AML cancer cell inhibition (Figure 1G). These results highlight the strong link between FTO enzymatic inhibition and the measured inhibitory effect in AML cells. Beyond their FTO inhibitory activity, RNB-557 and RNB-637 also exhibited favorable absorption, distribution, metabolism, and excretion (ADME) profiles (Figure 1H). Taken together, we obtained and optimized potent and selective FTO inhibitors, with high cellular activity, strong target engagement, and favorable pharmacological properties, positioning them as promising lead candidates for therapeutic development.

**Figure 1.**
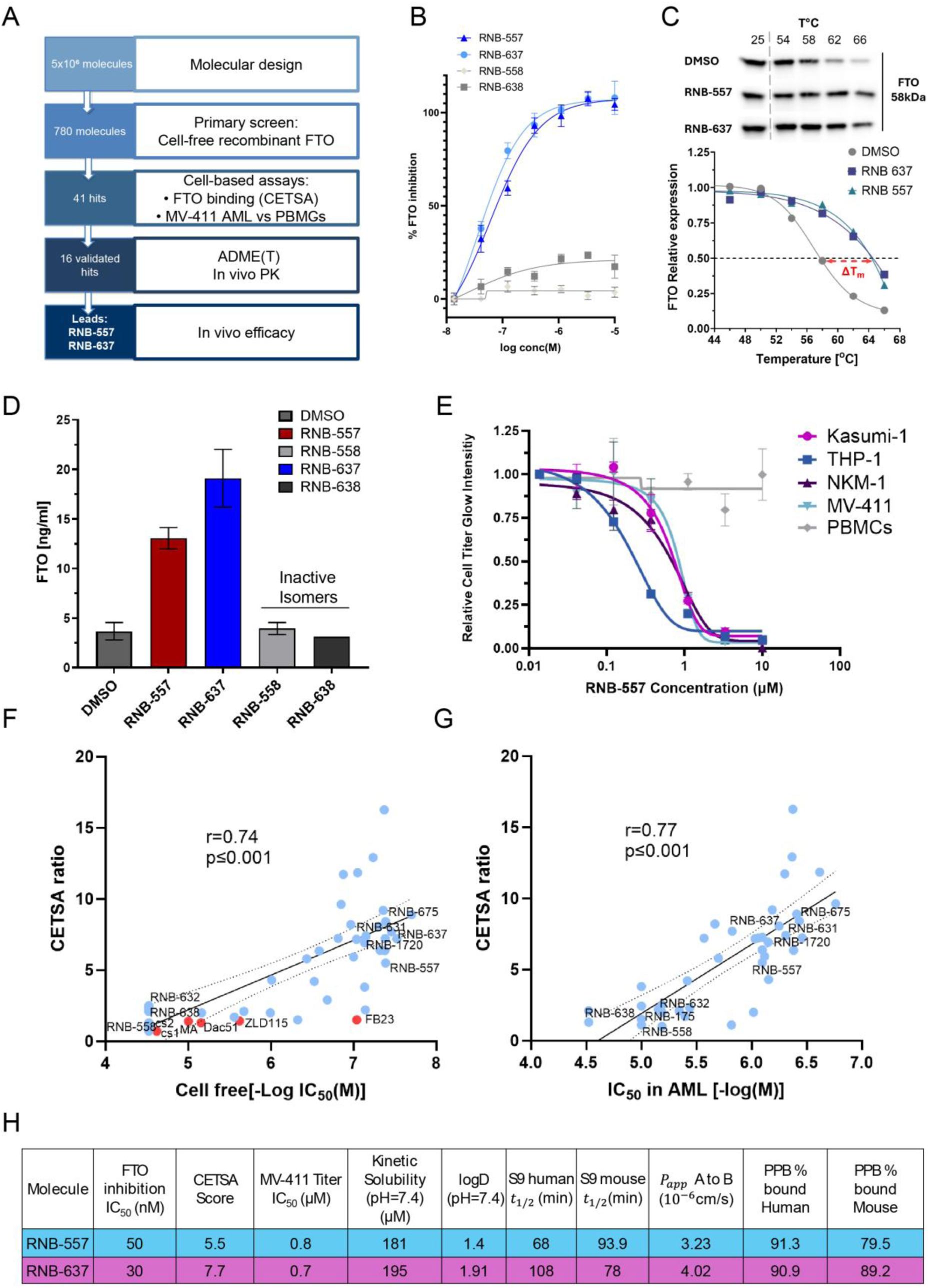
Drug discovery pipeline and early characterization of novel FTO inhibitors. **(A)** Schematic overview of the drug discovery workflow, from initial hit identification to the optimization phase that led to the selection of two lead candidates, RNB-557 and RNB-637. **(B)** FTO enzymatic inhibition assessed in a cell-free recombinant assay. Activity was normalized to DMSO (0% inhibition) and maximum signal (100%). **(C)** Cellular thermal shift assay (CETSA) performed in A2058 cells. Representative Western blot showing FTO stabilization in a temperature-dependent manner upon exposure to RNB-557, RNB-637 and DMSO negative control. At the bottom graph, signal quantification of FTO bands utilizing the ImageJ software. The signal was normalized to FTO levels at 25 °C. **(D)** CETSA quantification at a fixed denaturation temperature (62 °C) using competitive ELISA for measuring FTO protein amounts. Data represent mean ± SD from three experiments. **(E)** Dose-response cell titer after 96 hours exposure to RNB-557 across a panel of AML cell lines compared with PBMCs from a healthy donor. **(F)** CETSA binding scores (compound vs. DMSO at 62 °C) plotted against FTO cell-free half maximal inhibitory concentration (IC50) values (-logIC50(M)). Blue dots represent RNAble molecules derived from RNB-557/637. Red dots indicate known published FTO inhibitors. **(G)** CETSA binding scores (compound vs. DMSO at 62 °C) plotted against half-maximal effective concentration (EC50) in MV-411 AML cell line measured by Cell Titer Glow assay post treatment at the indicated concentrations. Cell titer at the lowest concentration was set to 1 and relative levels are shown. Values are mean ± SD from three different experiments. **(H)** RNB-557 and RNB-637 FTO inhibition, CETSA score, EC50 in MV-411 and ADME properties summary.

### FTO inhibition results in chromosome misalignment, prometaphase arrest and apoptosis in cancer cells

To gain deeper insight into the mechanism of cancer cell cytotoxicity induced by our hit series, we performed RNA-Seq analysis on RNA extracted from THP-1 AML cells 24 hours after treatment with 1 µM RNB-637 (Figure 2). Surprisingly, a set of mitotic genes including CCNB1 (Encoding for Cyclin B1 protein) were found to be significantly upregulated upon RNB-637 treatment while a set of DNA replication and G1/S transition genes, including CCNE1 (Encoding for Cyclin E1 protein) were downregulated (Figure 2A,B and Supplementary Figure S2A). Cyclin B1 protein is upregulated during late G2 and reaches its peak in mitosis, where it activates CDK1 to drive mitotic entry, rendering it a well-established marker of late G2 and mitotic phases^30^. RT-qPCR (Figure 2C) and immunofluorescence in BxPC-3 pancreatic cancer cells (Figure 2D,E) validated upregulation of CCNB1 pre-RNA, CCNB1-RNA and Cyclin B1 protein upon RNB-637 treatment. BxPC-3 cells were selected for subsequent imaging experiments based on their sensitivity to FTO inhibition (Figure 6D-F), adherent growth properties, and large cell size, which are particularly advantageous for microscopy analyses. Notably, RNB-637-treated BxPC-3 cells appeared to be arrested at mitosis (Figure 2D-DNA Enlarged). We then quantified DNA content within the cell population of DMSO control in comparison to 1 µM and 3 µM RNB-637 treatments and a clear, dose-dependent arrest at the G2/M phase was detected (Figure 2F). To gain deeper insight into the cell cycle arrest, we further monitored DNA and tubulin post RNB-637 treatment (Figure 2G,H). While all phases of mitosis were detected post DMSO control treatment, only prophase and prometaphase, were detected after RNB-637 treatment (Figure 2H), confirming a specific prometaphase mitotic arrest. In the RNB-637-treated prometaphase-arrested cells, chromosomes appeared misaligned, and the microtubule-organizing centers (MTOCs, or centrosomes) displayed a star-like morphology (Figure 2G, Merge-Enlarged). Chromosome mis-segregation and prolonged mitosis are tightly linked to activation of intrinsic stress pathways that frequently culminate in apoptosis^15^; therefore, we evaluated apoptosis following treatment with RNB-557 and RNB-637. As expected, both compounds induced a dose-dependent increase in apoptosis after 24 hours in THP-1 AML FTO sensitive cells, whereas the negative control isomer RNB-558 had no apoptotic effect (Supplementary Figure S2B,C).

**Figure 2.**
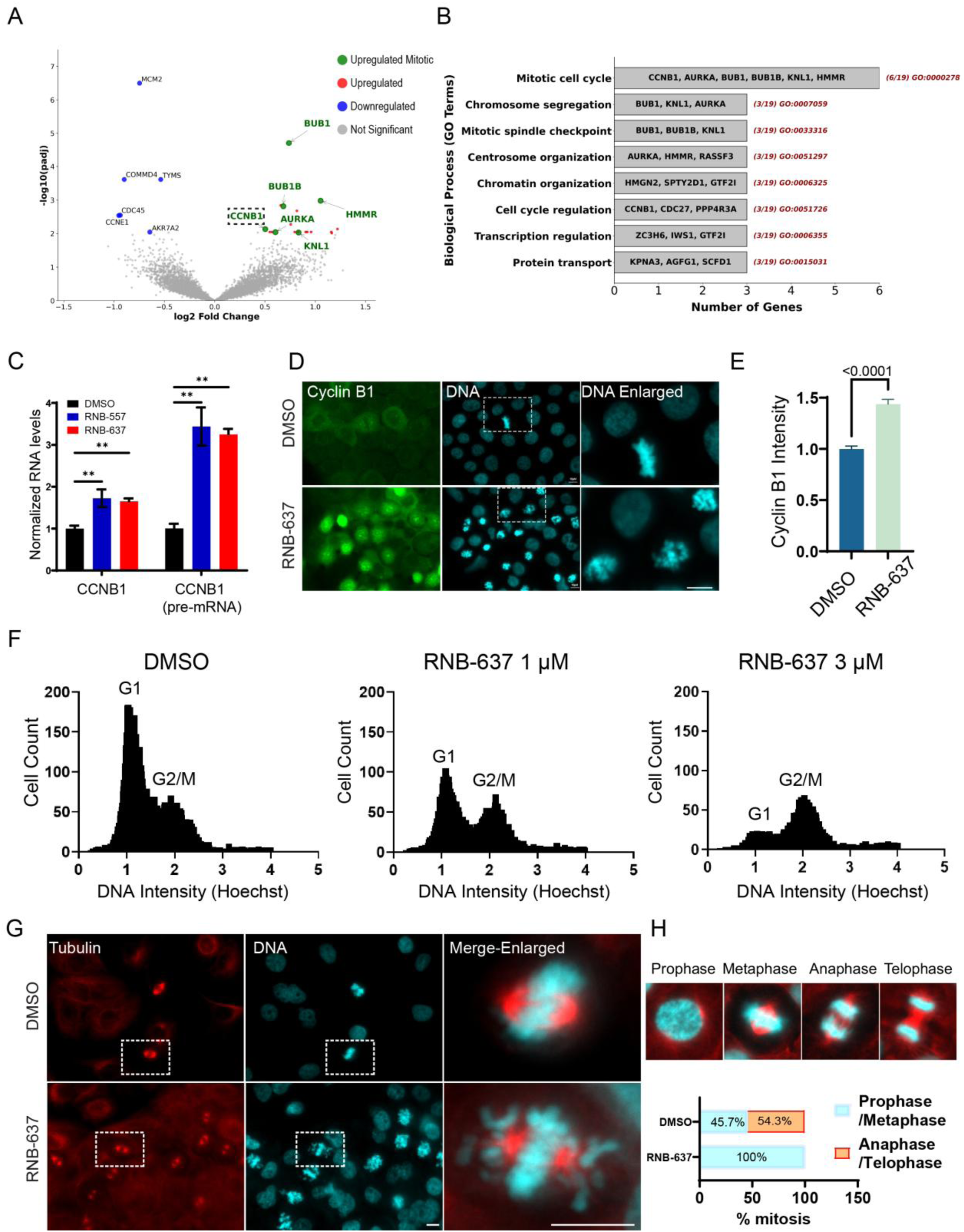
RNB-637 induces Cyclin B1 upregulation, prometaphase arrest and chromosomes misalignment. **(A)** RNA-sequencing analysis of THP-1 RNA purified after 24 hours treatment with RNB-637 (1 µM) versus DMSO control. Upregulated genes (-log(Padj) > 2, Log2(Fold Change) >0.5) are shown in red, mitotic upregulated genes in green and downregulated genes (-log(Padj) > 2, Log2(Fold Change) <-0.5) are shown in blue. Genes with no significant gene expression alteration (-log(Padj) < 2) are labeled in gray. **(B)** Gene ontology enrichment of the 19 significantly elevated genes. **(C)** RNA was purified from RNB-557 and RNB-637 BxPC-3 treated cells, (3 µM, 24 hours) followed by cDNA synthesis and RT-qPCR gene expression analysis with primers amplifying CCNB1 mRNA and CCNB1 pre-mRNA. Data represent mean ± SD from three experiments (n=3). Significance was determined by unpaired t-test (**, p-value < 0.01). **(D)** Immunofluorescence staining of CyclinB1 protein in RNB-637 (1 µM, 24 hours) and DMSO treated BxPC-3 cells. **(E)** Quantification of CyclinB1 protein immunofluorescence signal (integrated intensity in the nucleus). No primary control signal was subtracted, and values were normalized to DMSO control. n = 10 fields; unpaired t-test, p < 0.0001. **(F)** Quantification of Hoechst DNA staining of BxPC-3 cells following treatment with RNB-637 (1 µM and 3 µM, 24 hours). Values represent single cell integrated intensities. **(G)** Immunofluorescence of α-tubulin following 14 hours treatment with RNB-637 (3 µM) and DMSO. **(H)** Representative images of all the mitotic phases detected in DMSO control. Bottom panel, quantification of the specific mitotic phase detected after DMSO and RNB-637 (3 µM, 14 hours). Values represent mitotic events normalized to the total mitotic cell counts for each treatment (total of 25 mitotic cells per condition). (All scale bars = 10µm).

### FTO localizes to centrosomes and is essential for their structure and function during mitosis

Chromosome misalignment during mitosis can stem from defects in kinetochores, centrosomes, microtubule attachments, or specific motor protein activity^31^. To monitor chromosome misalignment during mitosis, we labeled the Aurora kinase B protein, known to localize at centromeres^32^. Following RNB-637 treatment, Aurora kinase B remained localized on individual chromosomes, suggesting intact kinetochore structures (Figure 3A), while FTO inhibition by RNB-637 induced a dose-dependent chromosome misalignment (Figure 3A,B). We then monitored FTO localization using immunofluorescence staining with an antibody validated for FTO specificity (Supplementary Figure S3A–D). Quantification of FTO in single cells revealed a clear dose-dependent elevation of FTO in RNB-637-treated cells, further demonstrating on-target binding of RNB-637 to cellular FTO (Supplementary Figure S3E,F). Additionally, FTO imaging revealed a distinct pool of FTO localized at centrosomes, as evidenced by colocalization with the centrosome marker pericentrin and tubulin across all tested human and mouse cell lines (Figure 3C-E, Supplementary Figure S3G,H) and during both interphase and metaphase (Figure 3E). Taken together, these findings suggest that FTO localizes to centrosomes and plays an essential role in centrosome-mediated chromosome alignment during mitosis.

**Figure 3.**
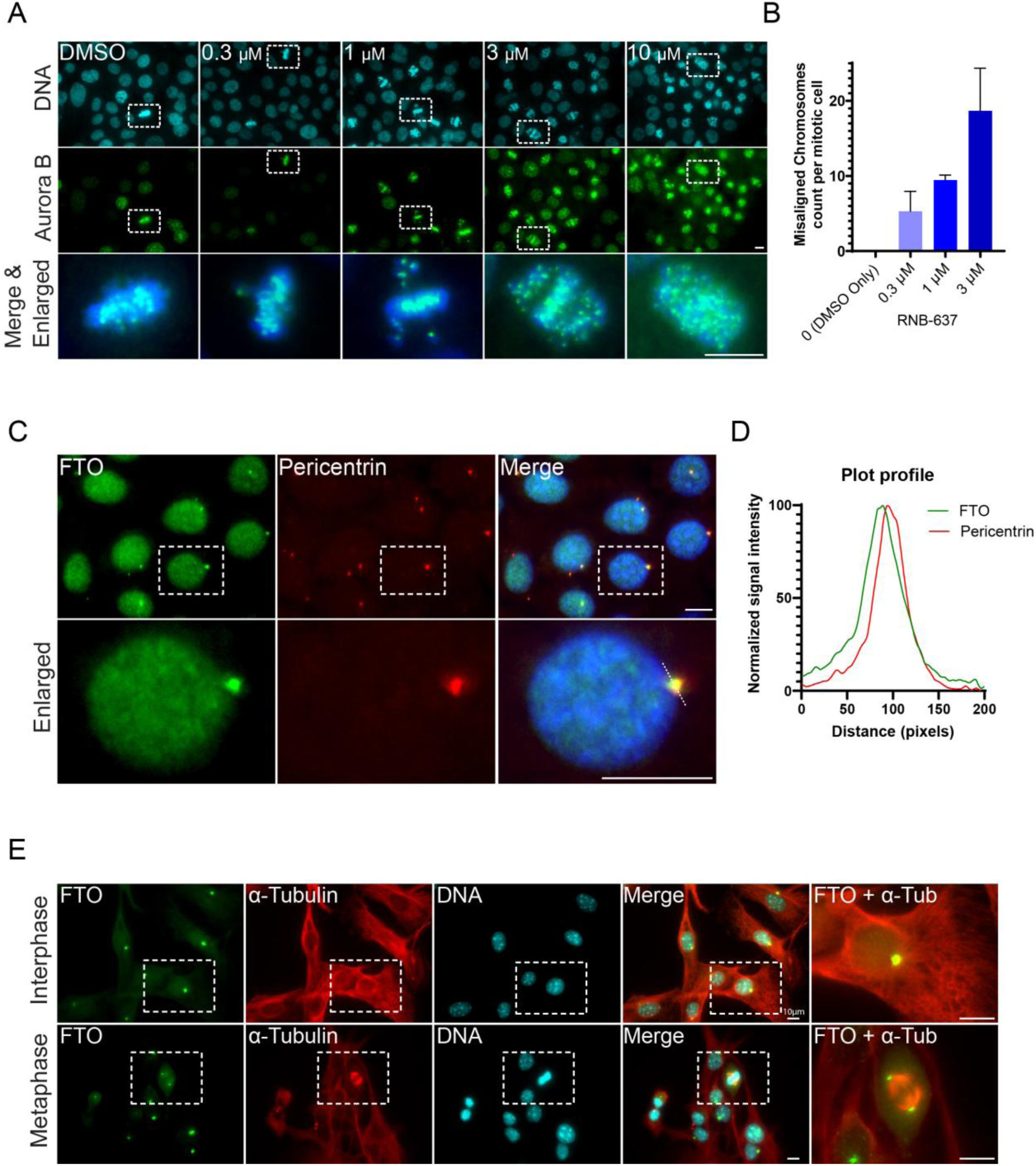
FTO localizes to centrosomes and its inhibition induces dose-dependent chromosome misalignment. **(A)** BxPC-3 cells treated for 14 hours with DMSO or RNB-637 in the indicated concentrations followed by immunofluorescence staining for Aurora kinase B protein and Hoechst DNA staining. Bottom panel, Merge&Enlarged, highlight representative mitotic cells (labeled by the white squares). **(B)** Quantification of chromosome misalignment across RNB-637 concentrations. Misalignment was scored based on deviation of single-counted chromosomes from the metaphase plate. At least ten mitotic cells were manually counted per condition; data represent mean ± SD. **(C)** Immunofluorescence co-staining of FTO and the centrosomal marker Pericentrin in BxPC-3 cells. Images captured at 60× magnification; scale bar, 10 µm. **(D)** Intensity profile analysis across centrosome across the white line shown in Merge/Enlarged. Intensities were background-subtracted and normalized to the maximum value within each data set. FTO(green) and Pericentrin(red) channels are shown. Image analysis was done utilizing ImageJ. **(E)** Immunofluorescence of α-tubulin and FTO in B16F10 murine melanoma cells, highlighting FTO localization at centrosomes. Enlarged insets show an interphase cell and a mitotic cell, with FTO enriched at the centrosomes and mitotic spindles, respectively. 60× magnification; scale bar, 10 µm.

### FTO inhibition leads to mitotic slippage

Since chromosome mis-segregation can lead to mitotic slippage^33^ as well as apoptosis (Supplementary Figure S2B), we evaluated the occurrence of mitotic slippage following treatment with RNB-557 and RNB-637. Indeed, in both BxPC-3 pancreatic cancer cells and OVCAR-3 ovarian cancer cells, which are also sensitive to FTO inhibition by RNB-637 (Figure 6F), an elevation of distinct population with >4N DNA content emerged after 48 hours of RNB-637 treatment (Figure 4A,B). These cells exhibited two or more distinct centrosome foci (Figure 4C-E), indicating that they had undergone mitotic slippage^34^. A similar phenotype was observed following FTO siRNA knockdown on both BxPC-3 and OVCAR-3 cells. Despite intermediate knockdown efficiency leaving some FTO at centrosomes (Supplementary Figure S4A,B), the proportion of cells undergoing mitotic slippage significantly increased (Figure 4F,G). These data, together with the centrosomal localization of FTO (Figure 3), support its role in centrosome-mediated chromosome alignment, which is essential for proper mitosis, and inhibition of FTO leads to either apoptosis or mitotic slippage.

**Figure 4.**
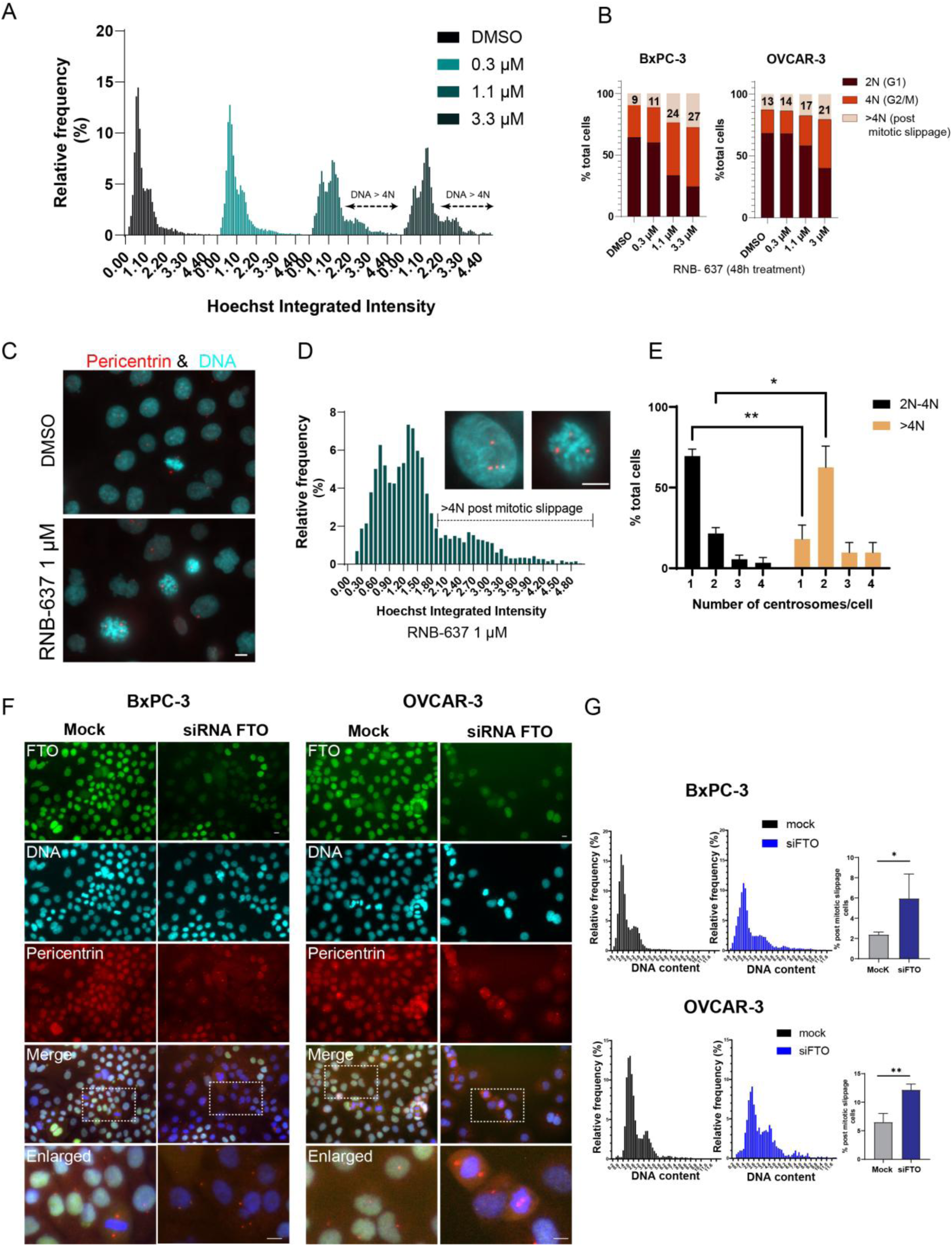
FTO inhibition triggers mitotic slippage via both pharmacological and genetic suppression. **(A)** Quantification of Hoechst DNA staining of single BxPC-3 cells following treatment with RNB-637 for 48 hours with the indicated concentrations. Arrows indicate emergence of post mitotic slippage cells exhibiting whole-genome duplication (>4N DNA content). Relative frequency values represent the relative cell count normalized to the total cell count in each treatment. **(B)** Quantification of relative BxPC-3 and OVCAR-3 cell count according to their DNA levels (2N, 4N and >4N) after 48 hours of DMSO or RNB-637 treatment at the indicated concentrations. Percentages of relative post-slippage (>4N) cells are indicated. **(C)** Immunofluorescence of Pericentrin (red) and DNA staining (cerulean) in BxPC-3 cells treated with DMSO and RNB-637 (1 µM, 48 hours) showing multipolar mitosis and large post-slippage cells harboring >2 centrosomes. 60× magnification, Max Projected Z stack is shown. scale bar, 10 µm. **(D)** Representative DNA intensity histogram demonstrating mitotic slippage (>4N Hoechst) at 1 µM treated cells. Above the plot are typical post-slippage cells (>4N) displaying aberrant centrosome numbers (Pericentrin, red). **(E)** Centrosome number distribution within 2N–4N versus >4N post-slippage single cells after 1 µM RNB-637, 14 hours treatment. Total of 90 cells were manually counted for the 2N-4N cell population, and 20 cells were counted for the >4N population. *, T-test p-value < 0.05. **, T-test p-value < 0.01. (**F)** FTO (green) and pericentrin (red) Immunofluorescence staining within BxPC-3 and OVCAR-3 cells, 144 h after FTO siRNA transfection or mock control. Enlarged images of FTO siRNA-treated cells depict cells that underwent mitotic slippage (DNA content >4N and centrosome count ≥2), compared to mock-treated controls **(G)** DAPI integrated-intensity of single BxPC-3 and OVCAR-3 cells following FTO siRNA for 144 h or mock control. Right panel: percentage of post-slippage (>4N) cells per condition (mean ± SD, n = 3). *, T-test p-value < 0.05. **, T-test p-value < 0.01.

### FTO is essential for intact NuMA and KIFC1 localization during mitosis

FTO has been reported to interact with the mitotic spindle proteins NuMA and KIFC1^35^, supporting its localization to centrosomes (Figure 3). Both NuMA and KIFC1 are essential for intact localization of each other and for chromosome segregation during mitosis^25^. Following FTO inhibition with RNB-637, KIFC1 mislocalized from the mitotic spindle to a diffusive cytoplasmic pattern as opposed to DMSO-treated controls where KIFC1 was strongly enriched at the spindles (Figure 5A,B). While KIFC1 mislocalized following RNB-637 treatment, other mitotic kinesins, Eg5 and TPX2 remained enriched at the mitotic spindles (Supplementary Figure S5A). As KIFC1 is a motor protein that affects the positioning of centrosomes, we monitored the distance between centrosomes in DMSO control and RNB-637-treated cells and revealed that RNB-637 treatment shortened the distance between pericentrin-labeled centrosomes in the arrested mitotic cells (Figure 5C). Interestingly, upon compound washout and replacement with fresh medium (RNB-637 Wash), KIFC1 rapidly regained proper localization in mitotic cells, and the distance between the two centrosomes was rapidly restored (Figure 5D,E and Supplementary Figure S5A). The rescued cells successfully completed mitosis, as evidenced by the restoration to a normal cell cycle distribution within 8 hours (Figure 5F). In addition to KIFC1 mislocalization, NuMA protein also changed pattern following FTO inhibition by RNB-637 (Figure 5G). Under normal conditions, NuMA localizes mainly to centrosomes in mitotic cells, as observed in the DMSO control, however, RNB-637 treatment caused a reduction in centrosomic NuMA levels (Figure 5G,H) in parallel to the appearance of foci throughout the cytoplasm (Figure 5G,I). RNB-637 wash eliminated these foci and restored NuMA centrosomal localization, along with proper KIFC1 positioning and chromatin alignment (Figure 5G–I). NuMA total amount did not change during either DMSO control, RNB-637 and RNB-637 wash, suggesting that FTO inhibition perturbs NuMA protein localization and not total expression (Figure 5G-I and Supplementary Figure S5B). Taken together, these results underscore the requirement of FTO activity for intact localization of the essential mitotic factors, NuMA and KIFC1. This does not exclude the possibility that FTO is also required for the localization of additional mitotic factors yet to be examined.

**Figure 5.**
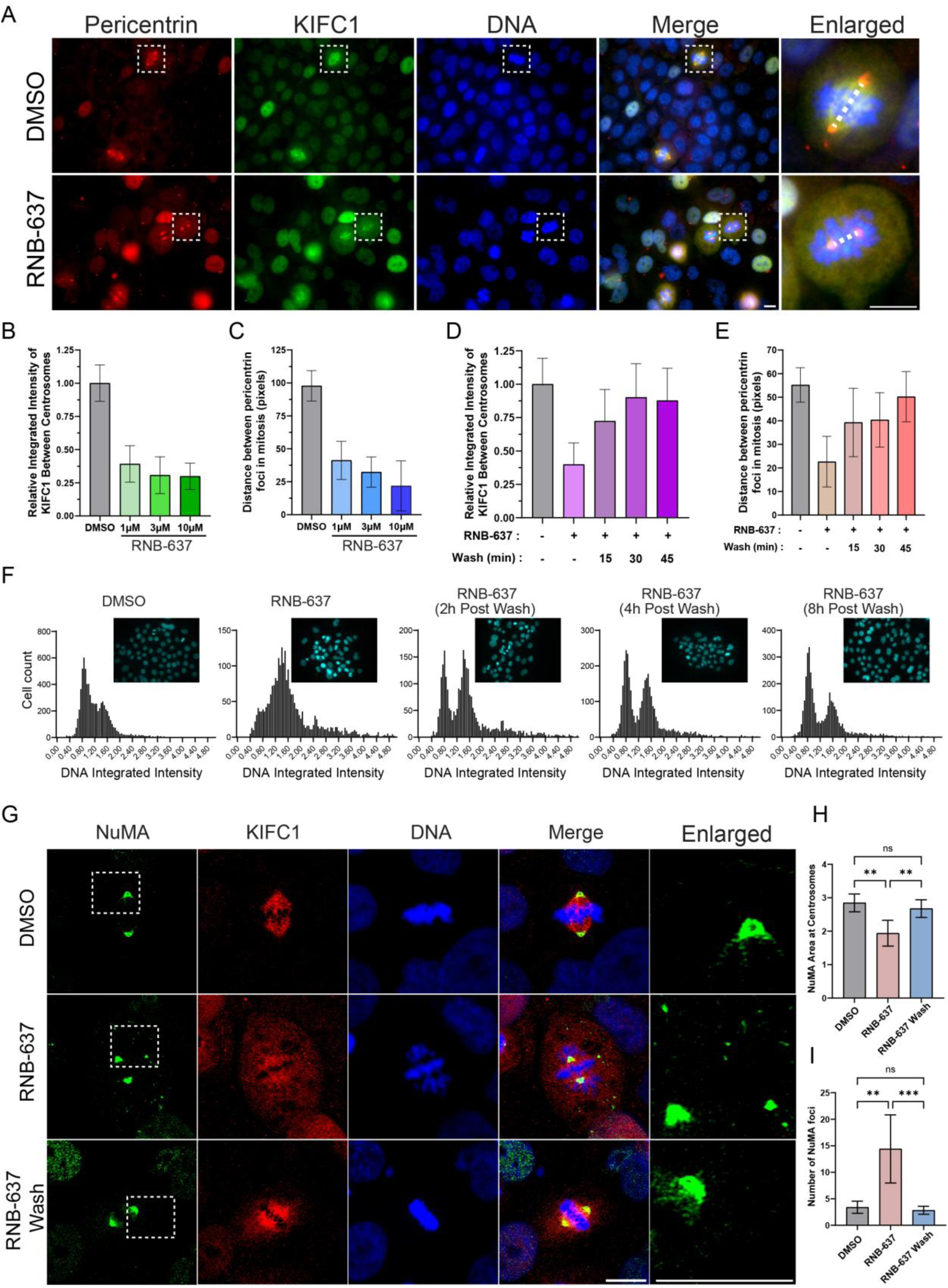
FTO inhibition induces mislocalization of KIFC1 and NuMA proteins during mitosis. **(A)** Immunofluorescence staining of KIFC1 (green) and Pericentrin (red) in BxPC-3 cells following 14 hours treatment with RNB-637 (3 µM). Enlarged images depict representative cells at mitosis in DMSO control and RNB-637 treated cells, shown in the white squares. 60X magnification, scale bar 10µm. **(B)** Quantification of KIFC1 fluorescence intensities between spindle poles after DMSO or RNB-637 at the indicated concentrations. **(C)** Quantification of the distance between poles in mitotic cells in the indicated treatments (quantification was made within at least 10 mitotic cells per treatment). **(D)** KIFC1 signal intensity between poles, following DMSO, RNB-637 (3 µM, 14 hours) or RNB-637 + replacement with media without RNB-637 (Wash) for the additional indicated times. (n = 10 fields). **(E)** Distance between poles measured following DMSO, RNB-637 (3 µM, 14 hours) or RNB-637 + replacement with media without RNB-637 (Wash) for the indicated times. (n = 10 fields, at least 10 mitotic cells per treatment). **(F)** Single cell DNA integrated-intensity distribution following DMSO and RNB-637 treatments (3 µM) for 14 hours as well as 14 hours following replacement with media without RNB-637 (Wash) for additional 2, 4, and 8 hours post replacement. Example fields, taken with a 40X magnification lens, are shown above, for each condition. **(G)** Immunofluorescence staining of NuMA, KIFC1 and DNA in DMSO after treatment with DMSO, RNB-637 (3 µM, 14 hours) and RNB-637 Wash (3 µM for 14 hours and then replacement with media without RNB-637 for additional 0.5 hours). Representative cells at mitosis are shown. (Z stack maximum projection, 60X). **(H)** Quantification of NuMA area at centrosomes in the same conditions described at G. (I) Quantification of NuMA foci count in mitotic cells in the same conditions as in G. In H and I, mitotic cells within five fields (n=5) were quantified per condition and the mean ± SD is shown.

### Cancer cells with CIN are specifically sensitive to FTO inhibition by RNB-637

As KIFC1 was previously reported to be specifically essential in cancers with centrosome amplification (CA), typically found in CIN cancers^24,36^, and FTO inhibition disrupted its proper localization in mitotic cells, we next tested whether cancer cells harboring CA and CIN are particularly sensitive to FTO inhibition. The ovarian cancer cell line OVCAR-3 was previously identified as harboring centrosome amplification^37^. As expected, centrosome counts in OVCAR-3 cells revealed that a substantial proportion of the cell population (>20%) contained more than one centrosome (CA+) (Figure 6A,B). In contrast, normal skin fibroblasts showed fewer than 10% of cells with more than one centrosome, likely reflecting the normal distribution across G1 and G2/M phases (Figure 6A,B). Similarly, we identified additional cell lines with centrosome amplification including BxPC-3 pancreatic and CAL 27 head and neck cancers. In contrast, U2OS osteosarcoma, 786-o renal cell carcinoma as well as normal fibroblasts did not show elevated centrosome amplification (Figure 6B). We further examined these cells for evidence of multipolar mitosis and found that CA+ cells were more likely to display this phenotype. Both features are recognized as indicators of chromosomal instability (CIN)^18^. Importantly, CIN+ cancer cells (CA+ and Multipolar mitosis+) showed greater sensitivity to FTO inhibition by RNB-637 compared with CIN- (CA- and Multipolar mitosis-) cells (Figure 6D-F). Overall, these data support a model in which FTO is essential for centrosome function and chromosome alignment during mitosis and that cancers with CIN, are particularly vulnerable to FTO inhibition. Taken together, these findings highlight FTO as a promising therapeutic target specific for these cancer types (Figure 6G).

**Figure 6.**
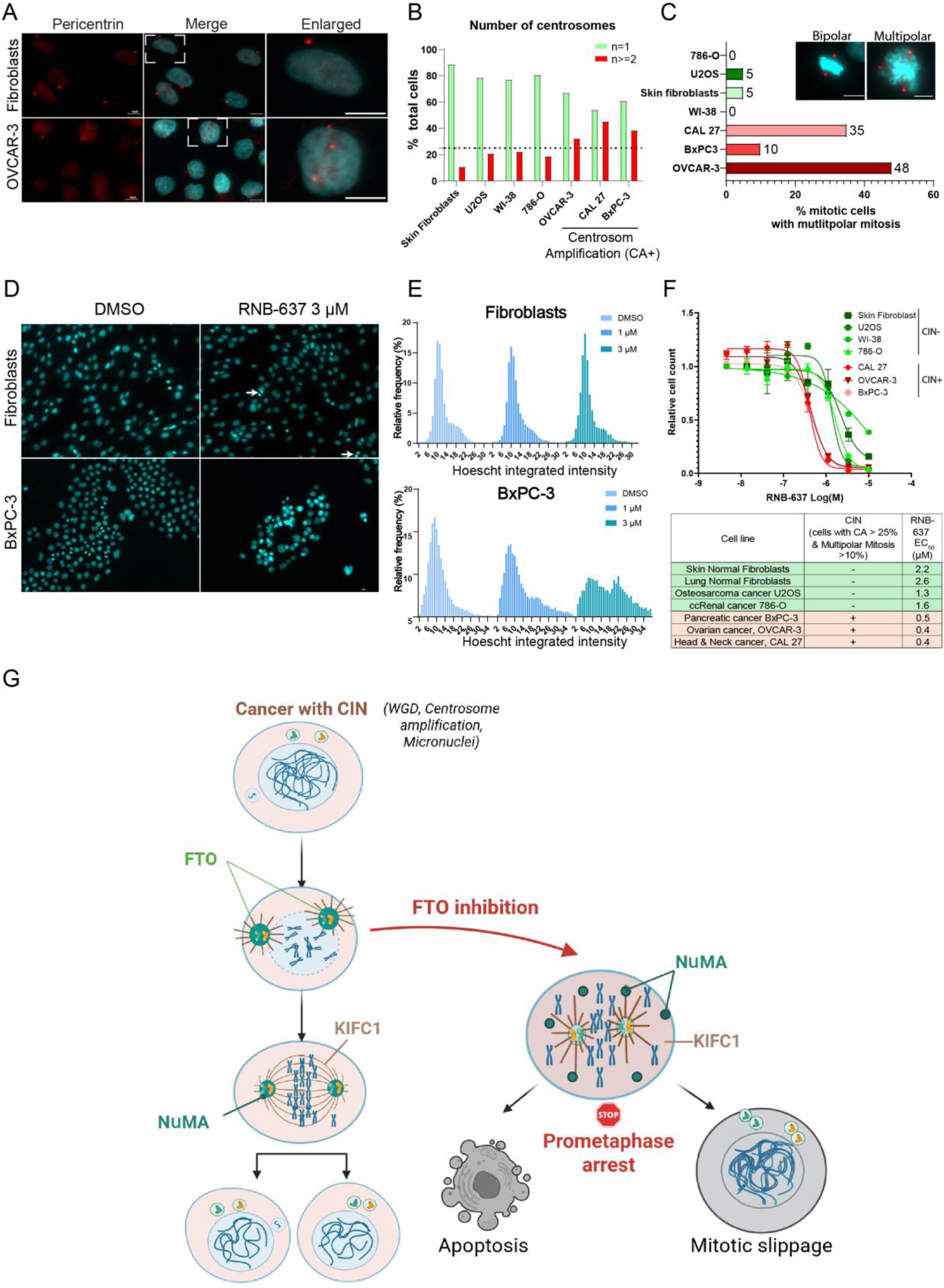
Chromosomal instability increases cancer sensitivity to FTO inhibition. **(A)** Immunofluorescence staining of Pericentrin and Hoechst DNA in skin fibroblasts and OVCAR-3 cells. Enlarged show representative examples of normal centrosome number in fibroblasts (one centrosome per cell) and centrosome amplification in OVCAR-3 (≥2 centrosomes). **(B)** Centrosome count per cell quantified across a panel of cell lines. Dashed line represents ≥25% cells with ≥2 centrosomes that is selected as cutoff for cancer with centrosome amplification (CA+). Minimum of 120 total cells were counted in 10 fields for each cell line. **(C)** Relative count of multipolar mitosis in the indicated cell lines (n = 20 mitotic cells per cell line). Images highlight representative normal bipolar and multipolar mitosis. **(D)** DNA Hoechst staining in BxPC-3 and skin fibroblasts treated with RNB-637 (3 µM, 14 hours). White arrows indicate cells at telophase, observed in skin fibroblasts but absent in BxPC-3 under identical treatment conditions. **(E)** Quantification of Hoechst DNA integrated-intensity within single BxPC-3 and normal skin fibroblasts treated with DMSO and RNB-637 at the indicated concentrations. **(F)** Relative cell counts, 96 hours post treatment with RNB-637 at the indicated concentrations (Log(M)) of the indicated cell types, labeled as CIN- and CIN+, according to the threshold set in B. Bottom Table summarizes the EC_50_ values and CIN status (CIN -, Green and CIN +, Red) for each cell line tested. **(G)** Mechanistic model: In CIN+ cancer cells, FTO inhibition induces NuMA and KIFC1 mislocaliztion, chromosome misalignment and mitotic arrest, leading to apoptosis or mitotic slippage. Created in BioRender. https://BioRender.com/tc1ch9v.

### In vivo efficacy of FTO inhibition by RNB-637

Previous studies and our ongoing research have shown that AML cell lines are sensitive to FTO inhibition (Figure 1E and Supplementary Figure S1E). Among tested AML cell lines, THP-1 cells displayed the highest sensitivity to RNB-637 (EC_50_ = 200 nM). These cells contain an enlarged centrosome compared to PBMCs from a healthy donor, which may point on the presence of two or more clustered centrosomes per centrosome foci (Supplementary Figure S6A,B). In addition, THP-1 cells contain high levels of multipolar mitosis (Supplementary Figure S6C), indicative of CIN. Additionally, THP-1 cells exhibited RNB-557/637-dependent elevation of mitotic cells and post-mitotic slippage, while these effects were not observed upon treatment with the inactive isomer control RNB-558 (Supplementary Figure S6D). Taken together, these results indicate that THP-1 cells are chromosomally unstable and sensitive to chromosome misalignment upon FTO inhibition by RNB-557 and RNB-637. We next aimed to evaluate the efficacy of RNB-637 against THP-1 AML in vivo. CB17 SCID mice were inoculated with THP-1 cancer cells, and tumor growth was monitored. When tumors reached 100 mm^3^, mice were randomly divided into four groups and were treated orally twice daily for 21 days with RNB-637 at 25, 50, or 75 mg/kg, or with vehicle control. RNB-637 demonstrated a clear dose-dependent antitumor effect: 25 mg/kg showed minimal activity (Tumor Growth Inhibition (TGI) = 14.1%), 50 mg/kg produced moderate inhibition (TGI = 56.7%), and 75 mg/kg achieved strong tumor suppression (TGI = 81.7%; Supplementary Figure S6E,F), with no detectable toxic effects. Further, pharmacokinetic (PK) analysis revealed a clear exposure-efficacy relationship, with serum concentrations at the 75 mg/kg dose substantially exceeding the THP-1 half-maximal effective concentration (EC_50_) throughout the dosing interval (Supplementary Figure S6G). To extend these findings beyond hematological malignancies and evaluate the efficacy of RNB-637 in a solid tumor model, we next tested an optimized RNB-637 lead in OVCAR-3 (CIN+, Figure 6A-C) ovarian cancer xenografts. BALB/c nude mice were inoculated subcutaneously with OVCAR-3 cells, and treatment was initiated when tumors reached 100 mm³. Mice were randomly divided into four groups and treated orally twice daily for 33 days at 25, 50, or 75 mg/kg, or with vehicle control. A clear dose-dependent antitumor effect was observed: 25 mg/kg produced modest inhibition (TGI = 32.5%), 50 mg/kg achieved moderate tumor suppression (TGI = 44.3%), and 75 mg/kg resulted in strong antitumor activity (TGI = 90.5%), with no detectable toxic effects (Figure 7A,B). Tumor weights measured on the final day corroborated the volumetric data, further confirming dose-dependent efficacy (Figure 7C). PK analysis revealed a clear exposure-efficacy relationship, with serum concentrations at the 75 mg/kg dose substantially exceeding the OVCAR-3 EC_50_ throughout the dosing interval (Figure 7D). Collectively, these findings demonstrate that our novel FTO inhibitors exhibit potent antitumor activity in both hematological malignancies and solid tumor models characterized by chromosomal instability, supporting their continued development as promising therapeutic candidates.

**Figure 7.**
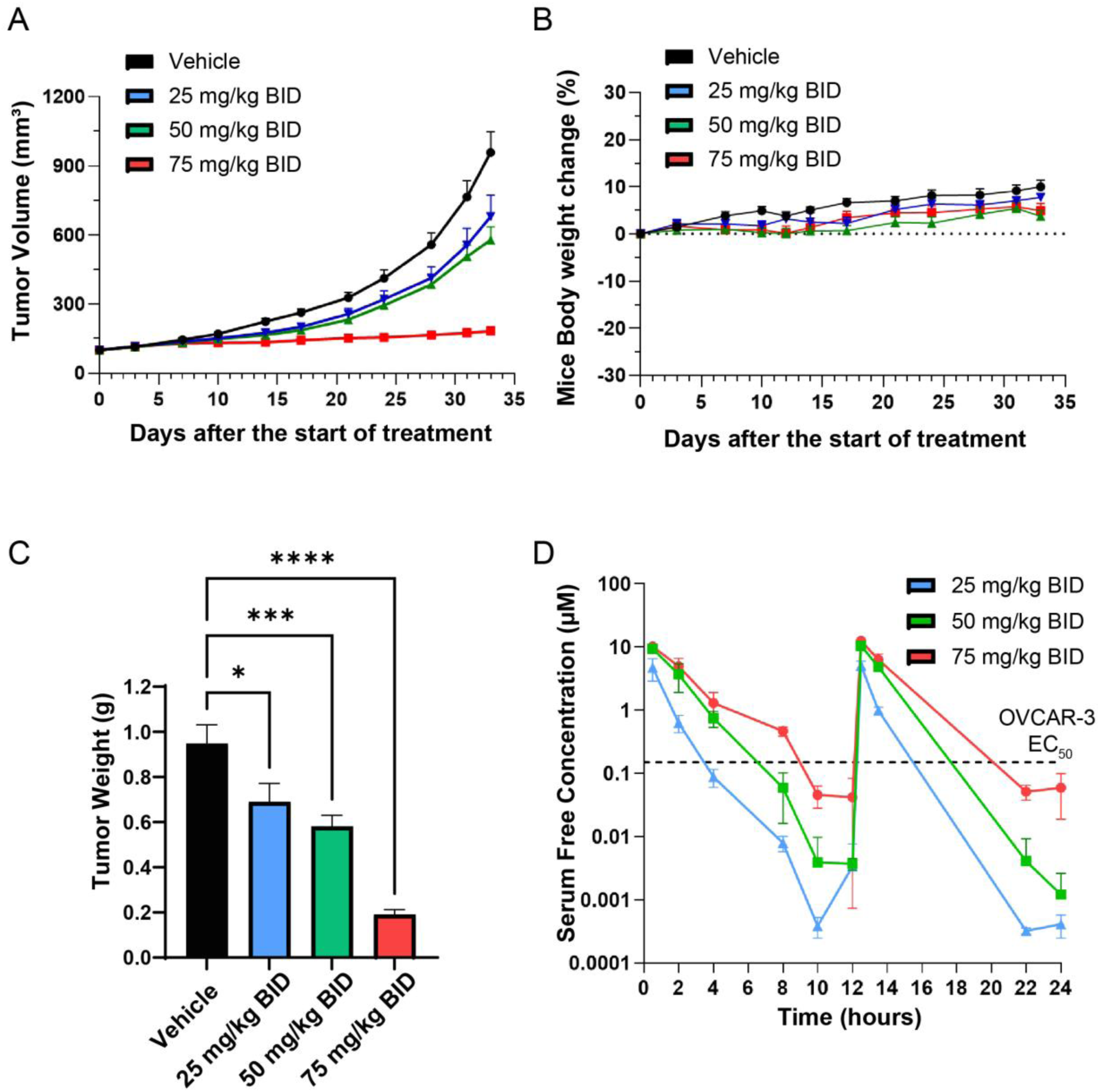
CIN-high ovarian solid tumor exhibit selective vulnerability to FTO inhibition in-vivo. **(A)** In vivo OVCAR-3 xenografts in BALB/c nude mice treated with vehicle control and RNB-637 optimized analog dosed orally, twice per day, at 25, 50, and 75 mg/kg or vehicle control, for 33 days. Mean tumor volume ± SEM is shown (n = 10 mice/group). **(B)** Mice body weight throughout the experiment duration. Mean mice weight ± SEM is shown (n = 10 mice/group). **(C)** On day 33, tumors were excised, and the mean weight ± SEM is shown. Two-way Anova p-value is shown (p-value < 0.05 (*), p-value < 0.001 (***), p-value < 0.0001(****)). **(D)** Pharmacokinetic analysis of RNB-637 optimized analog concentrations following oral administration in BALB/c nude mice. Blood samples were collected from three mice per treatment group at multiple time points over 24 hours on day 33. Mean serum free compound concentration ± SEM is shown. The dashed line indicates the OVCAR-3 EC50 value determined in vitro.

## Discussion

Our findings establish FTO as a critical regulator of centrosome function and chromosome segregation during mitosis. To our knowledge, this study provides the first evidence of centrosomal localization of FTO, a feature conserved across mice and human cells (Fig. 3C,E). While FTO contains a canonical nuclear localization signal (NLS) and predominantly localizes to the nucleus, previous studies have reported a cytoplasmic fraction of the protein^38^. In line with this expanded subcellular distribution, the centrosome-associated proteins NuMA and KIFC1 were previously identified in the proximal proteome of endogenous FTO using BAR SILAC–MS, revealing an unexpected spatial association between FTO and the centrosome machinery^35^.

By showing that FTO localizes to centrosomes and that its inhibition mislocalizes key centrosomal proteins such as NuMA and KIFC1, we provide direct mechanistic evidence linking FTO to the assembly of intact spindles during mitosis. These perturbations further led to apoptosis or mitotic slippage, known to be mutually exclusive, dependent on the specific cell^39^. As FTO is an RNA demethylase, we demonstrated that RNB-637 inhibits FTO m6A-RNA demethylase activity (Figure 1B). Therefore, we predict that FTO function in centrosome-mediated spindle assembly is linked to RNA metabolism. Several studies had shown that RNA transcripts localized to centrosomes, including NUMA1 and CCNB1 mRNAs^40–42^. Specifically, NUMA1 mRNA localization to centrosomes was shown to be dependent on active translation, which initiates at the cytoplasm and then translocates to the centrosomes with the nascent translated protein^42^. Recently, FTO was shown to be essential for localized translation of stress transcripts^43^. Specifically, it was shown that upon stress, FTO translocates to the cytoplasm and undergoes phosphorylation by the stress response kinase MARK4. Upon activation, FTO demethylates HARP transcripts, thereby increasing their translation and contributing to cancer cell survival. The translation is localized on distinct γ-Tubulin and FTO-associated ribosomes. Thus, FTO might regulate specific γ-Tubulin RNA translation during stress and γ-Tubulin/centrosomic RNA translation during mitosis.

In addition to RNA localization to centrosomes, a recent study reported on RNA-dependent protein-protein interactions during mitosis^44^. Specifically, the authors identified KIFC1 as a mitotic protein whose interaction with other mitotic proteins, Aurora Kinase A and TPX2, is dependent on RNA. Given that we observed mislocalization of both KIFC1 and NuMA proteins upon FTO inhibition, future studies should aim to elucidate the specific RNA processing events regulated by FTO that coordinate proper localization of NuMA and KIFC1 proteins during mitosis.

An important implication of our work is the selective vulnerability of cancers harboring CIN to FTO inhibition. This selective sensitivity was evident across multiple cell lines and extended to in vivo ovarian and AML xenograft models, in which RNB-637 and RNB-637 optimized analog produced marked and dose-dependent tumor growth inhibition through 21 days (Supplementary Figure S6E) and 33 days (Figure 7) without detectable toxicity. Errors originating during mitosis, including chromosome alignment, can lead to chromosomal instability (CIN) which is indicated by the presence of cells with centrosome amplification (CA) and multipolar mitosis within the tumor population^45^. Specifically, KIFC1 has been reported to be essential for cancers with high centrosome amplification^24,36^, which aligns with our findings. KIF18A was also identified as an important target in CIN cancers^20–23^. Future studies should determine whether FTO regulates KIF18A RNA and protein localization during mitosis, as well as elucidate the full spectrum of FTO-mediated RNA processing events critical for mitotic progression.

In summary, our study identifies FTO as a centrosomal protein essential for accurate chromosome segregation during mitosis, a process particularly vulnerable in CIN cancers, thereby providing a strong mechanistic rationale for targeting FTO and positioning the selective FTO inhibitor, RNB-637 optimized lead, as a promising candidate for clinical development in these tumor types.

## Methods

### Cell culture

#### Adherent cells

Human cell lines, U2OS (osteosarcoma), WI-38 (lung fibroblasts), A549 (lung adenocarcinoma), BxPC-3 (pancreatic ductal adenocarcinoma), OVCAR-3 (ovarian carcinoma), CAL-27 (tongue squamous cell carcinoma), 786-O (renal cell carcinoma), A2058 (melanoma), human skin fibroblasts and the murine melanoma cell line B16F10 were cultured as adherent monolayers. U2OS, A549, BxPC-3, CAL-27, 786-O, and A2058 cells were cultured in Dulbecco’s Modified Eagle Medium (high glucose DMEM, Sartorius #01-055-1A) supplemented with 10% fetal bovine serum (FBS, Gibco, catalog # A5256701) and 1% penicillin–streptomycin (Merck, catalog #P4333). WI-38 cells were maintained in Eagle’s Minimum Essential Medium (EMEM, Sartorius, #01-025-1A) supplemented with 10% FBS, 1% penicillin–streptomycin, 1× non-essential amino acids (Merck, #M7145), and 1 mM sodium pyruvate (Sigma-Aldrich, #S8636). OVCAR-3 cells were cultured in RPMI-1640 medium (Sartorius, #01-106-1A) supplemented with 20% FBS, 1% penicillin–streptomycin, 0.01 mg/mL bovine insulin (Merck #11070-73-8), and 1 mM sodium pyruvate, in accordance with ATCC recommendations. Healthy skin fibroblasts (GM05659) were obtained from the Coriell Institute for Medical Research and originated from a healthy male donor (1 year old at the time of sampling). Unless otherwise stated, all cell lines were obtained from the American Type Culture Collection (ATCC) and maintained according to ATCC recommendations at 37°C in a humidified incubator with 5% CO₂. Cells were routinely tested and confirmed to be mycoplasma negative. All adherent cells were passaged using 0.25% trypsin–EDTA (IM Beit HaEmek #L0931) before reaching full confluency and were used within a limited number of passages (<25) to ensure phenotypic stability.

#### Suspension cells

Human leukemia cell lines Kasumi-1, MV4-11, and THP-1 were cultured in suspension in RPMI-1640 medium supplemented with 10% FBS and 1% penicillin–streptomycin. Cells were maintained at densities recommended by ATCC and passaged by dilution with fresh medium.

#### PBMCs isolation

Peripheral blood mononuclear cells (PBMCs) were isolated from whole blood of healthy donors by density-gradient centrifugation using Ficoll-Paque PLUS (Cytiva, #17144002) according to the manufacturer’s instructions. PBMCs were cultured in RPMI-1640 supplemented with 10% FBS and 1% penicillin–streptomycin and used within a limited number of days after isolation.

#### Chemistry

All compounds were designed in-house and synthesized by Piramal Pharma Ltd and WuXi Apptec Co., Ltd. Compound purity was ensured by ^1^H-NMR, LCMS and HPLC measurements and exceeded 95% purity for all tested compounds. Batches used for animal testing have a purity of >98%.

#### RNA extraction

Cells were seeded in 6-well plates and allowed to adhere overnight prior to treatment. Following treatment, cells were lysed, and total RNA was extracted using the ISOLATE II RNA Mini Kit (Bioline, #BIO-52072) according to the manufacturer’s instructions, including an on-column DNase I treatment for 10 minutes to eliminate residual genomic DNA. RNA concentration and purity were assessed using a NanoDrop spectrophotometer by measuring absorbance at 260 nm.

#### RNA sequencing

Total RNA extracted as described above was used for RNA-sequencing analysis. Integrity of the isolated RNA was tested using the Agilent High Sensitivity RNA Kit and Tapestation 4200 at the Genome Technology center at the Faculty of Medicine Bar-Ilan University. 100 ng of total RNA were used for libraries preparation using the Quantseq 3’ mRNA-Seq prep kit FWD (Lexogen). Quantification of the library was performed using dsDNA HS Assay Kit and Qubit 2.0 (Molecular Probes, Life Technologies) and qualification was done using the Agilent D1000 Tapestation Kit and Tapestation 4200. 50nM of each library was pooled together and was diluted to 4nM according to NextSeq manufacturer’s instructions. 1.55pM was loaded onto the Flow Cell with 1% PhiX library control. Libraries were sequenced on an Illumina NextSeq 500 instrument, 75 cycles single read sequencing. RNA-seq analysis was performed with three biological replicates per condition. Each sample yielded between 2.0 and 6.3 million reads. FastQ Screen^46^ was used to exclude potential contamination from ribosomal, bacterial, viral, or other non-human sources. Prior to alignment, the first 12 nucleotides of each read were trimmed. Filtered reads were aligned to the human reference genome (GRCh38, annotation release 111) using STAR^47^. Gene-level read counts were generated using featureCounts (v1.6.0)^48^, and normalization and differential expression analyses were performed using DESeq2 (v1.42.1)^49^.

#### Reverse transcription and quantitative PCR (RT-qPCR)

Complementary DNA (cDNA) was synthesized from total RNA using the LunaScript® RT SuperMix Kit (New England Biolabs, #E3010) following the manufacturer’s instructions. Quantitative PCR (qPCR) was performed using SYBR™ Green Universal Master Mix (Applied Biosystems™, #4309155) together with gene-specific forward and reverse primers diluted to 10 µM (see Supplementary Information: Primers, siRNA, RNA Oligos, and Antibodies). No-reverse-transcription (No-RT) controls were included in each experiment to verify the absence of DNA contamination. Relative gene expression was calculated using the comparative Ct (2^(⁻ΔΔCq)) method, with HPRT1 or GAPDH serving as endogenous reference genes (see Supplementary Information: Primers, siRNA, RNA Oligos, and Antibodies).

#### Immunofluorescence and DNA staining

Cells were seeded in 96-well plates (Greiner Bio-one, #655182) or on 13 mm #1 cover glass (Bar Naor, #BN1001-13-1CN) prior to treatment. After treatment, the medium was removed, and cells were fixed with 4% paraformaldehyde (PFA; Santa Cruz, #sc-281692) for 20 minutes. Following three washes with 1× phosphate-buffered saline (PBS; Merck, #D1408), cells were permeabilized with 0.1–0.5% Triton X-100 (Sigma-Aldrich, #T8787) for 10 minutes. Blocking was performed using 5% bovine serum albumin (BSA; Merck, #810533) for 30 minutes at room temperature. Cells were then incubated overnight at 4 °C with primary antibodies (see Supplementary Information: Primers, siRNA, RNA Oligos, and Antibodies). The next day, cells were washed three times with PBS and incubated for 1 hour at room temperature with appropriate secondary antibodies (see Supplementary Information). After three additional PBS washes, Hoechst 33342 (Merck, #14533) was added at a concentration of 10 µg/mL for 20 minutes, followed by a final PBS wash before imaging. For DNA staining only (without immunostaining), Hoechst staining was performed immediately after Triton permeabilization.

### Imaging and Image analysis

#### Cell count and DNA histograms

After Hoechst staining, images were acquired using a Thermo Fisher EVOS FL automated microscope. Autofocus was applied to each well based on the Hoechst signal. Images were captured with a 4× objective lens and analyzed using a custom CellProfiler pipeline^50^ (v4.0.7) employing the Adaptive Otsu method to identify individual nuclei. Cell counts and the frequency distribution of integrated Hoechst signal intensities were generated and plotted using GraphPad Prism (v10.6.1).

#### Centrosome amplification quantification

Cells were seeded on coverslips and stained for Pericentrin and Hoechst as described above. Images were acquired using a 60× oil immersion objective and analyzed with a custom CellProfiler (v4.0.7) pipeline. The pipeline was configured to detect nuclei, extrapolate the cytoplasmic area, and generate a mask of individual cells. Within this mask, centrosomes were identified using the EnhanceSpeckles module, and each centrosome was assigned to its corresponding cell using the RelateObjects module. The number of centrosomes per cell was quantified and further analyzed and plotted as histograms for each cell line using GraphPad Prism (v10.6.1).

#### Distance between centrosomes and NuMA & KIFC1 quantification

After Pericentrin and NuMA or KIFC1 immunostaining (see above), individual mitotic cells were selected for analysis using ImageJ (v1.54p). A line was drawn between the centers of the two centrosome foci, and both the distance and the fluorescence intensity profile across this line for NuMA or KIFC1 were quantified. For NuMA foci quantification, foci were detected using the ImageJ^51^ Analyze Particles tool. Data were further analyzed and plotted using GraphPad Prism.

#### RNA interference and transfections

Cells were seeded 14 hours prior to transfection. siRNA (see Supplementary Information: Primers, siRNA, RNA Oligos, and Antibodies) transfection was performed using Lipofectamine™ RNAiMAX Transfection Reagent (Invitrogen, Cat#13778075) according to the manufacturer’s instructions. Following the specified incubation period and treatments, cells were either fixed for imaging, harvested for RNA purification or for protein extraction.

#### CETSA

Cells were collected and resuspended in PBS (1×) at a density of 1 × 10⁷ cells/mL. The suspension was mixed with the tested compound to achieve a final concentration of 30 µM, 0.3% DMSO in a 96-well PCR plate. Cells were incubated for 1 hour at 37 °C, followed by thermal treatment for 3 minutes at either a single temperature or across a temperature gradient using a StepOnePlus™ Real-Time PCR system. Samples were then lysed by three freeze–thaw cycles (liquid nitrogen followed by room temperature), followed by centrifugation at 17,000 × g for 20 minutes at 4 °C to pellet cell debris. The resulting supernatant was transferred to a new collecting tube for further FTO protein detection and quantification by western blot or competitive ELISA (see below).

#### Western blot

Except for CETSA experiments (see above), in all other experiments, the treated cells were lysed using RIPA buffer (Cat# 89900, Thermo) supplemented with cOmplete™ Protease Inhibitor Cocktail (Cat# 04693116001, Roche). Briefly, cells were washed with PBS (1×) and incubated with 200 µL of lysis buffer for 20 minutes on ice. Lysates were collected and centrifuged at 17,000 × g for 20 minutes at 4 °C to remove cell debris. All protein samples were prepared by dilution with XT Sample Buffer (4×; Cat#1610791, Bio-Rad) and supplemented with XT Reducing Agent (20×; Cat#1610792, Bio-Rad). Gel electrophoresis was performed using 4–12% Criterion™ XT Bis-Tris Protein Gel (Cat#3450125, Bio-Rad) with XT MOPS Running Buffer (Cat#1610788, Bio-Rad). Proteins were transferred to a nitrocellulose membrane (BioRad Trans-Blot Turbo 0.2µm Nitrocellulose, #1704158 and 1704159) using a semi-dry transfer system (BioRad Trans-Blot Turbo Transfer System). After transfer, membranes were blocked in blocking buffer; 5% non-fat dry milk (Merck, #70166) in Tris-buffered saline (Holland Moran, #J60877.1) with 0.1% Tween-20 (Sigma-Aldrich, #P6585)(TBST) for 1 hour at room temperature to prevent nonspecific binding. Membranes were then incubated overnight at 4 °C with the indicated primary antibodies (see Supplementary Information: Primers, siRNA, RNA Oligos, and Antibodies) diluted in blocking buffer. The following day, membranes were washed three times with TBST and incubated for one hour at room temperature with HRP-conjugated secondary antibodies. After additional washes, protein bands were detected using enhanced chemiluminescence (ECL) reagents (Merck, #WBLUF0500) and visualized using a digital imaging system (Azur biosystems, Azur 400). Signal intensity was quantified using ImageJ Gels tool and data was further analyzed and plotted using GraphPad Prism.

#### Competitive ELISA for FTO protein quantification

High-binding Nunc MaxiSorp plates (Cat# 442404, Invitrogen) were coated with recombinant FTO protein at a concentration of 10 ng/mL (expressed and purified by Prof. Reuven Wiener’s laboratory, Hebrew University of Jerusalem) and incubated overnight at 4 °C. Cell lysates obtained after thermal treatment or recombinant FTO standards for calibration were pre-incubated with anti-FTO antibody (17 ng/mL; Abcam, #ab126605) overnight at 4 °C. Recombinant FTO standards were serially diluted to generate a standard curve ranging from 50 ng/mL to 0.78 ng/mL. Pre-coated plates were washed three times with phosphate-buffered saline containing 0.05% Tween-20 (PBST) and blocked for 1 hour at room temperature with PBS supplemented with 5% bovine serum albumin (BSA) to prevent nonspecific binding. After removing the blocking buffer, the pre-incubation mixture of anti-FTO antibody and lysates and FTO standards was added in triplicate to the coated plates and incubated for 2 hours at room temperature. Plates were then washed three times with PBST and incubated with horseradish peroxidase (HRP)-conjugated secondary antibody for 30 minutes at room temperature (see Supplementary Information: Primers, siRNA, RNA Oligos, and Antibodies). Following three additional washes with PBST, Pierce™ TMB Substrate solution (Pierce™, #TS-34021) was added, and the reaction was allowed to develop in the dark for 10–20 minutes. The enzymatic reaction was stopped by adding 1 M sulfuric acid, and absorbance was measured at 450 nm with a reference wavelength of 570 nm using a Tecan® Spark microplate reader.

#### Recombinant FTO enzymatic assay

Recombinant FTO protein was expressed and purified by Prof. Reuven Wiener’s laboratory (Hebrew University of Jerusalem). The enzymatic reaction buffer was freshly prepared and contained 238 µM ammonium iron (II) sulfate hexahydrate (Sigma-Aldrich, #203505), 300 µM α-ketoglutarate (Merck, #K1128), 2 mM ascorbic acid (Merck, #A0278), and 50 mM HEPES (Merck, #H0887). Tested compounds were first diluted in DMSO using 1:2 serial dilutions and then further diluted 1:300 in enzymatic buffer prior to incubation with FTO (0.3 µM) for 5 min at room temperature. Following pre-incubation, the FTO–compound mixtures were incubated with a biotinylated single-stranded m⁶A modified RNA substrate or an unmodified RNA control (see Supplementary Information: Primers, siRNA, RNA Oligos, and Antibodies) at a final concentration of 50nM for 30 min at room temperature. After the reaction, FTO was heat-inactivated for 5 min at 95 °C, and samples were subsequently kept on ice. For m⁶A-RNA detection by ELISA, reactions were loaded onto streptavidin-coated 96-well plates (Greiner, #655990) and incubated overnight at 4 °C to allow binding. Plates were washed three times with PBST (0.05% Tween-20) and incubated for 1 hour at room temperature with an anti-m⁶A antibody (see Supplementary Information). Plates were washed three additional times and incubated with HRP-conjugated anti-rabbit secondary antibody for 30 min at room temperature in 5% BSA. Following three washes, TMB substrate (Pierce, #34021) was added, and the reaction was allowed to develop for 15 minutes. The reaction was stopped with 2 M sulfuric acid, and absorbance was measured at 450 nm using a Tecan® Spark microplate reader. Each plate included the following controls: DMSO + FTO, ssRNA substrate alone, unmethylated oligo RNA substrate (background control), and FTO incubated with a known inhibitor. IC₅₀ values were calculated from dose–response curves after normalization to the maximum signal (ssRNA substrate alone) and DMSO control (set to 0). Data were analyzed and plotted using GraphPad Prism.

#### Cell titer

Cells were seeded into 96-well plates and allowed to adhere overnight. The following day, cells were treated with the indicated concentrations of test compounds, with the final concentration of DMSO maintained at 0.1% (v/v) in all conditions. At the end of the treatment period, cell viability was assessed using the CellTiter-Glo® Luminescent Cell Viability Assay (Promega, #G7570) according to the manufacturer’s instructions. Luminescence was measured using a Tecan® Spark microplate reader. Background signal from cell-free medium was subtracted, and luminescence values were normalized to the DMSO-treated control. Data were analyzed and plotted using GraphPad Prism. For cell counts and titters by imaging see Methods/Cell count and DNA histograms.

#### Annexin V/Propidium Iodide Apoptosis Assay

Cells were seeded in 24-well plates and treated with the indicated concentrations of test compounds, maintaining a final DMSO concentration of 0.1% (v/v) in all conditions. After 24 hours of incubation, cells were harvested and stained using the Dead Cell Apoptosis Kit with Annexin V for Flow Cytometry (Invitrogen, Cat# V13241) according to the manufacturer’s instructions. Stained cells were analyzed on a MACSQuant® Analyzer 10 flow cytometer, and data were processed and visualized using FlowJo™ software (BD Biosciences).

#### In vivo efficacy

CB-17 SCID mice were inoculated subcutaneously into the right flank with THP-1 tumor cells (10 × 10⁶) suspended in 0.2 mL PBS supplemented with Matrigel (1:1 v/v). BALB/c nude mice were inoculated subcutaneously into the right flank with OVCAR-3 tumor cells (10 × 10⁶) in 0.2 mL PBS supplemented with Matrigel (1:1 v/v). Treatment was initiated when mean tumor volume reached approximately 100 mm³. Animals were assigned to treatment groups by stratified randomization based on tumor volume using Excel-based randomization software, with 10 tumor-bearing mice per group. The test article was formulated in 5% DMSO, 5% Solutol, and 90% water and administered orally twice daily at the indicated doses. Tumor dimensions were measured twice weekly using calipers, and tumor volume was calculated using the formula V = 0.5 × a × b², where a and b represent the longest and shortest tumor diameters, respectively. Body weight and general health status were monitored throughout the duration of the experiments. On the final day of the experiment, three mice per treatment group were selected for pharmacokinetic analysis, after which all tumors were excised and weighed.

## Acknowledgments

The authors acknowledge the use of AI-based tools, including Claude 3.5 Sonnet (Anthropic, 2024) and ChatGPT (accessed via Microsoft Copilot in Microsoft Word), for editorial assistance in text refinement and language editing. Figure 2A and Supplementary Figure S2A (volcano plot and GO terms analysis) were generated with assistance from Claude 3.5 Sonnet (Anthropic, 2024).

---

**Supplementary Figure S1.**
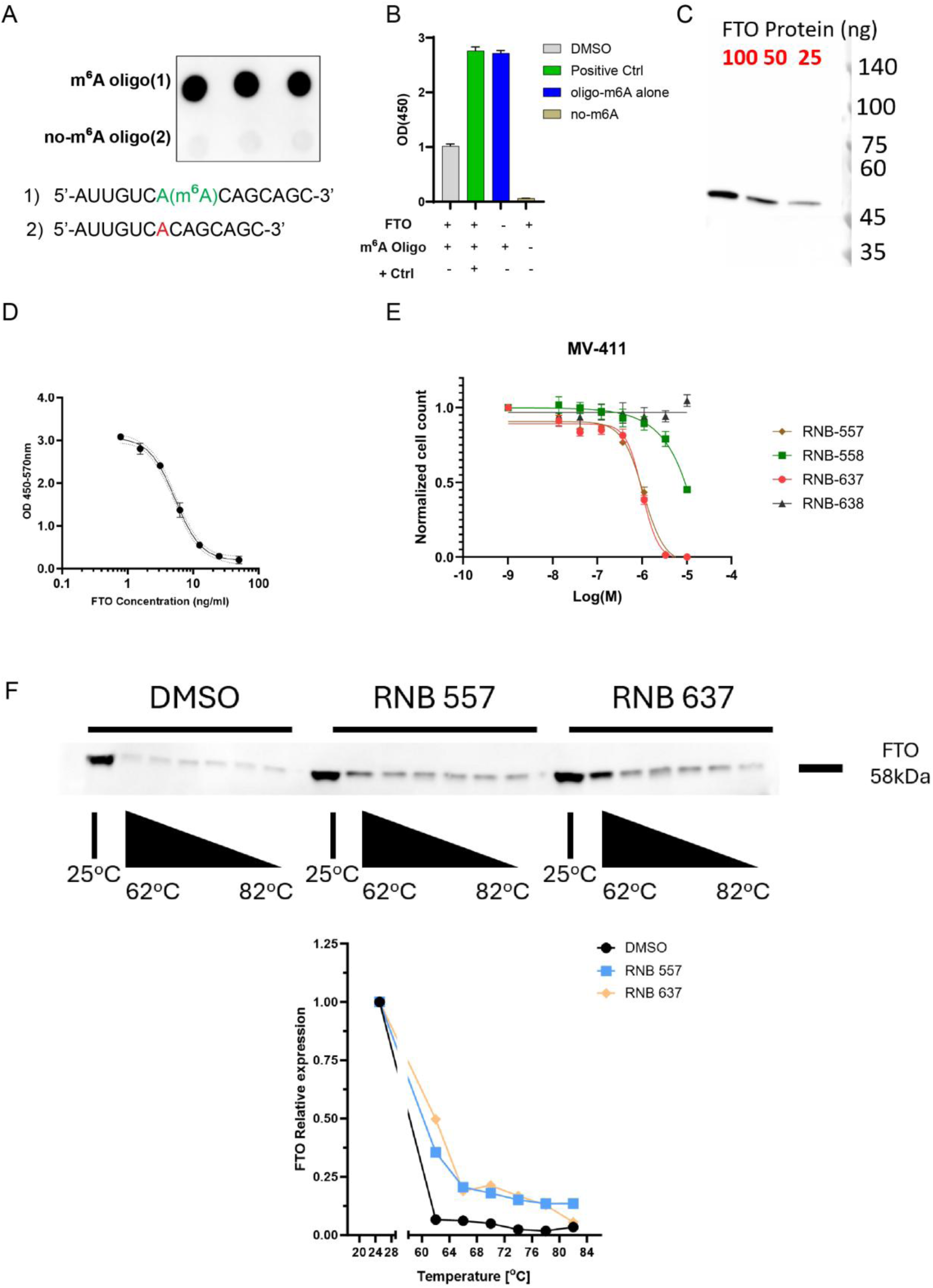
**(A)** m⁶A dot blot using RNA oligonucleotides containing m⁶A modification (1, m⁶A Oligo) or no m⁶A modification (2, no-m⁶A Oligo). **(B)** Recombinant FTO cell-free enzymatic reaction with 1-m⁶A Oligo (m⁶A Oligo +) or 2-no-m⁶A Oligo (m⁶A Oligo -) shown in A, with or without the FTO enzyme and with or without FB23, 10 µM, that was used as an FTO inhibitor positive control (+ Ctrl). m⁶A-RNA signal was detected by ELISA with an anti-m⁶A -RNA primary antibody (see supplementary information). Optical density at wavelength of 450nm (OD (450)), which is proportional to the amount of m⁶A -RNA is shown. Values within +FTO and + Oligo-m6A were set to 1 and the relative mean ± SD (n = 3 replicates) is shown. **(C)** Western blot detection of the recombinant FTO protein that was utilized in the cell free assay shown in B. The same antibody was used for FTO detection in cells and cell lysates generated during CETSA studies shown in D. **(D)** Standard curve generated with recombinant FTO protein for competitive FTO ELISA for measuring FTO amounts that was served as the end point measurement in high throughput CETSA assay. **(E)** MV-411 AML cell titter after 96 hours treatment with RNB-557, RNB-637 or their inactive isomers, RNB-558 and RNB-638. Cell titers at the lowest concentrations were set to 1 and relative cell titers are shown for each compound at the different concentrations. Mean ± SEM from three technical repeats is shown (n=3). **(F)** CETSA performed in mouse B16-F10 cells. Representative western blot showing FTO stabilization in a temperature-dependent manner upon exposure to RNB-557 (30 µM, 1 hour) RNB-637 (30 µM, 1 hour) and DMSO (0.1%, 1 hour) are shown. At the bottom graph, signal quantification of FTO bands utilizing the ImageJ software. The signal was normalized to FTO levels at 25°C.

**Supplementary Figure S2.**
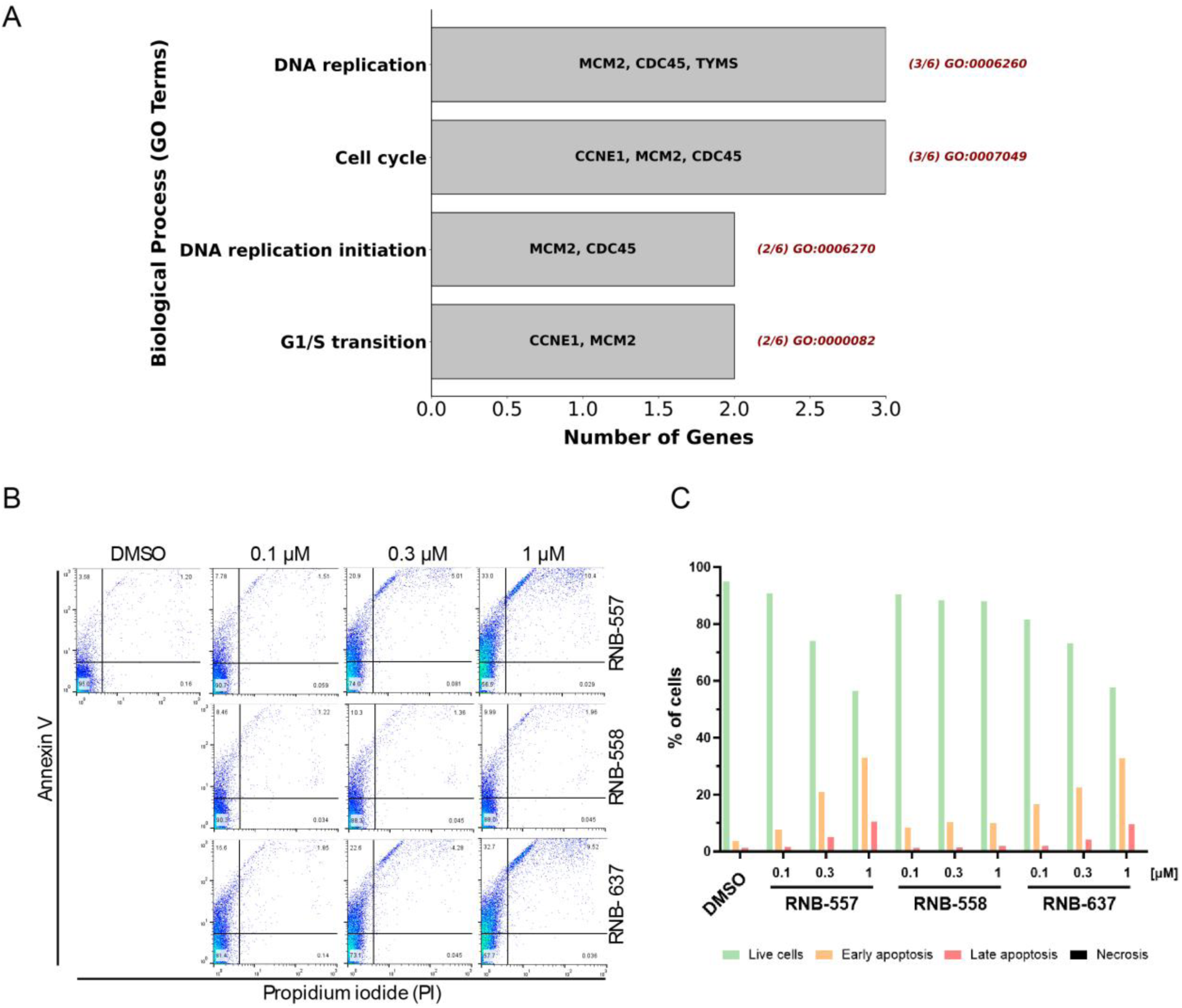
**(A)** RNA-sequencing analysis of THP-1 RNA purified after 24 hours treatment with RNB-637 (1µM) versus DMSO control. Gene ontology enrichment of the 6 significantly downregulates genes (-log(Padj) > 2, Log2(Fold Change) <-0.5) is shown. **(B)** Flow cytometry analysis of THP-1 cells stained with Annexin V/PI apoptosis detection kit. **(C)** Quantification of live cells, early apoptotic cells, late apoptotic cells and cells at necrosis following treatment with RNB-557, RNB-637, or the inactive analog, RNB-558. Relative cell counts normalized to the total cell number per condition is shown.

**Supplementary Figure S3.**
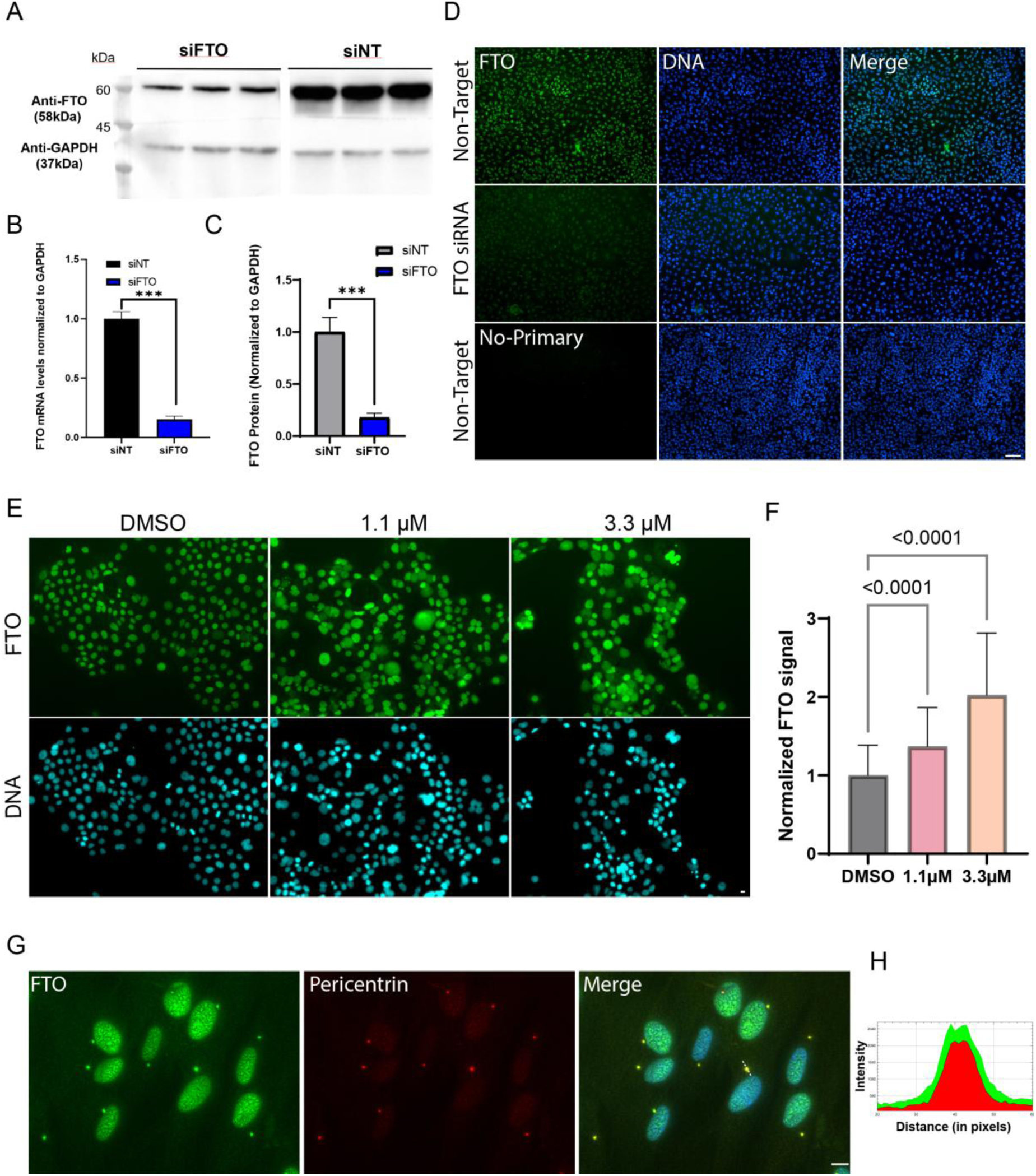
**(A)** Western blot analysis of FTO and GAPDH expression in A549 cells 72 h after siRNA-mediated FTO knockdown (n = 3 technical replicates). **(B)** RT-qPCR analysis of FTO mRNA expression normalized to GAPDH mRNA in A549 cells following siRNA-mediated knockdown vs non target siRNA (NT). Values of siNT were set to 1 and the mean + SEM (n = 3 technical replicates) relative expression is shown. ***, unpaired T test p-value < 0.001 **(C)** Quantification of FTO protein shown in A. Signal quantification of FTO bands was made by the ImageJ software. FTO signal was normalized to GAPDH and values of siNT were set to 1. Relative normalized FTO protein levels mean + SEM (n = 3 technical replicates) is shown. ***, unpaired T test p-value < 0.001**. (D)** Immunofluorescence staining of FTO (green) in A549 cells after siRNA-transfection of Non-Target or FTO siRNA. Cells were fixed 72 hours post siRNA transfection before staining. Scale bar, 100µM. **(E)** Immunofluorescence of FTO in BXPC-3 cells following 14 hours treatment with RNB-637. Images acquired utilizing a X4 magnification lens **(F)** Quantification of FTO immunofluorescence signal normalized to DMSO control. Image analysis was performed using a custom CellProfiler pipeline. Values represent integrated intensity mean + STD of cells within 3 different fields (DMSO-8123 cells, 1µM-4579 cells, 3 µM-3584 cells). Unpaired t-test P-values are shown. **(G)** Immunofluorescence of FTO protein in lung fibroblasts (60×), maximum Z-projection. **(H)** Plot profile of pixel intensity of FTO (green) and pericentrin (red) across a representative centrosome (dashed white line in Merge).

**Supplementary Figure S4.**
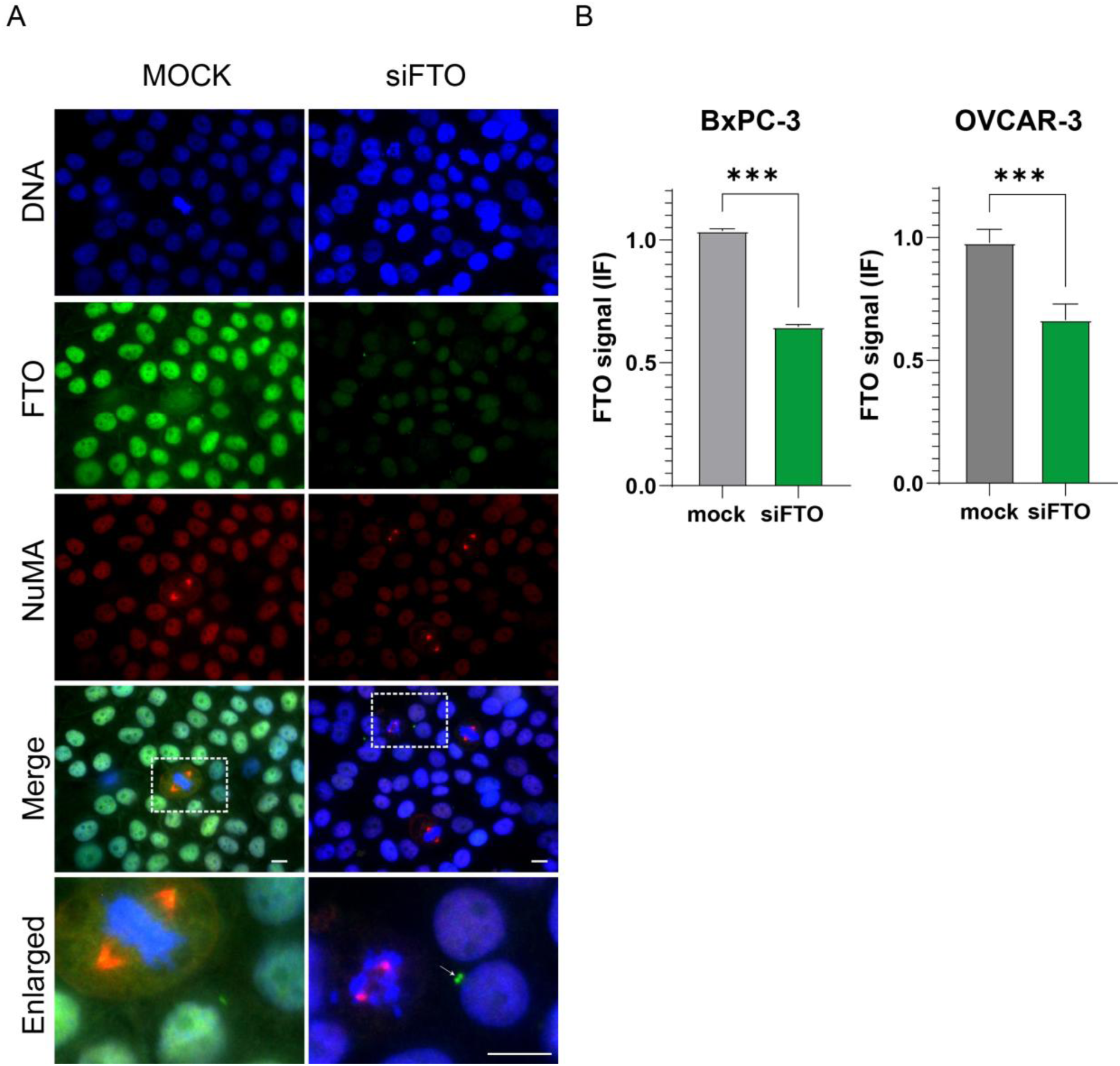
**(A)** Immunofluorescence staining of FTO (green) and NuMA (red) in BXPC-3 cells 144 h after transfection with FTO siRNA (siFTO) or Mock. X60 lens images are shown. Arrow at Enlarged show centrosomal localization of remaining FTO after siFTO treatment. **(B)** Quantification of FTO immunofluorescence signal following siRNA treatment in BxPC-3 and OVCAR-3 cells. Mean FTO intensity was normalized to mock control. Data represents relative FTO mean ± SD levels from 3 technical repeats (n = 3). ***, unpaired T test p-value < 0.001.

**Supplementary Figure S5.**
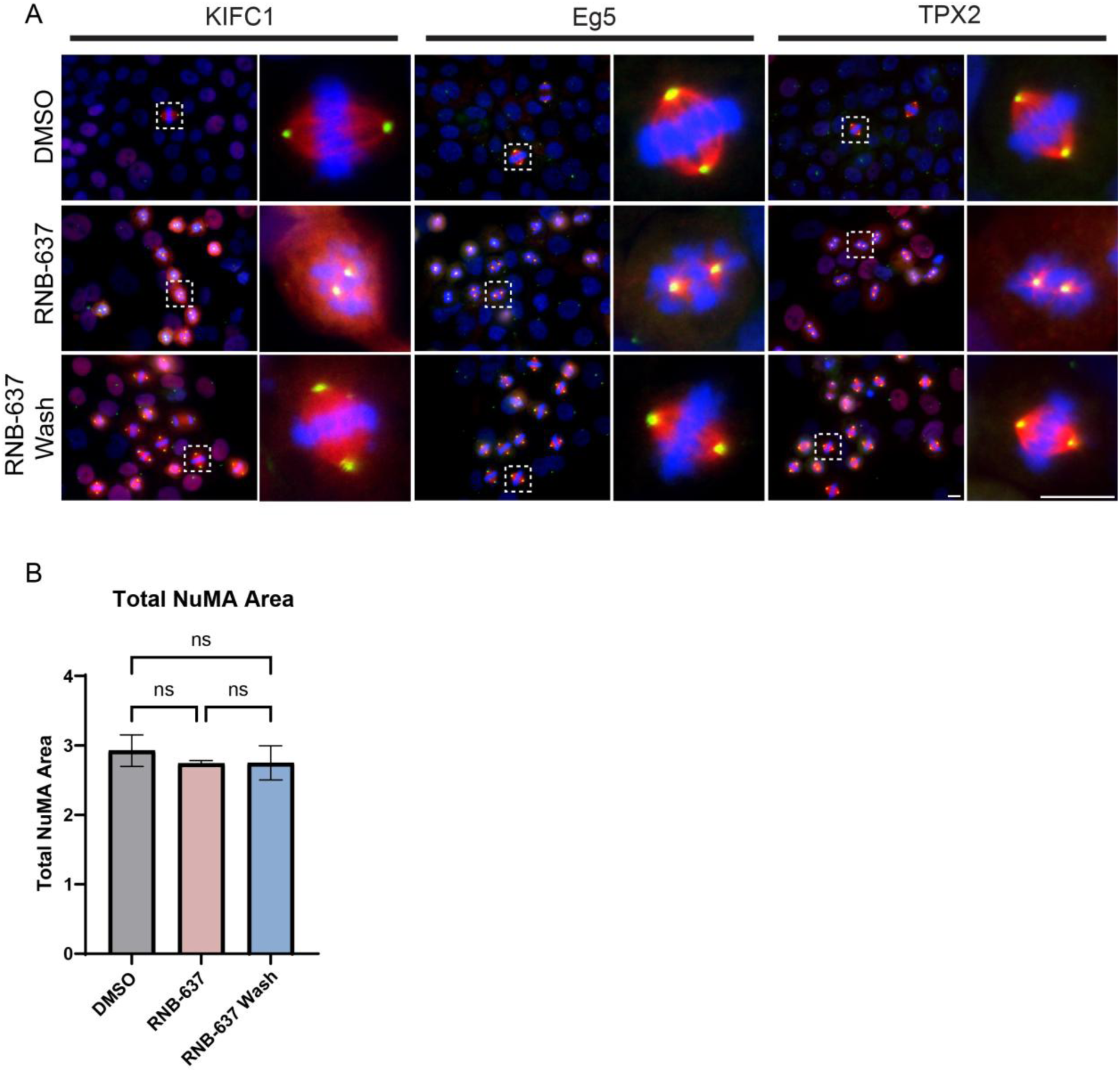
**(A)** Immunofluorescence staining of Pericentrin (green) and either KIFC1, Eg5, or TPX2 (red) in BxPC-3 cells. Cells were treated for 14 hours with DMSO or RNB-637 (3 µM), followed by either immediate fixation or compound washout and a 30-minute incubation in fresh medium prior to fixation (RNB-637 Wash). DNA was labeled by Hoechst staining (blue). **(B)** Quantification of total NuMA protein area, including centrosomes and NuMA foci, in BXPC-3 cells treated with DMSO, 3 µM RNB-637, or following RNB-637 Wash. Mitotic cells within five fields (n=5) were quantified per condition and the mean ± SD is shown.

**Supplementary Figure S6.**
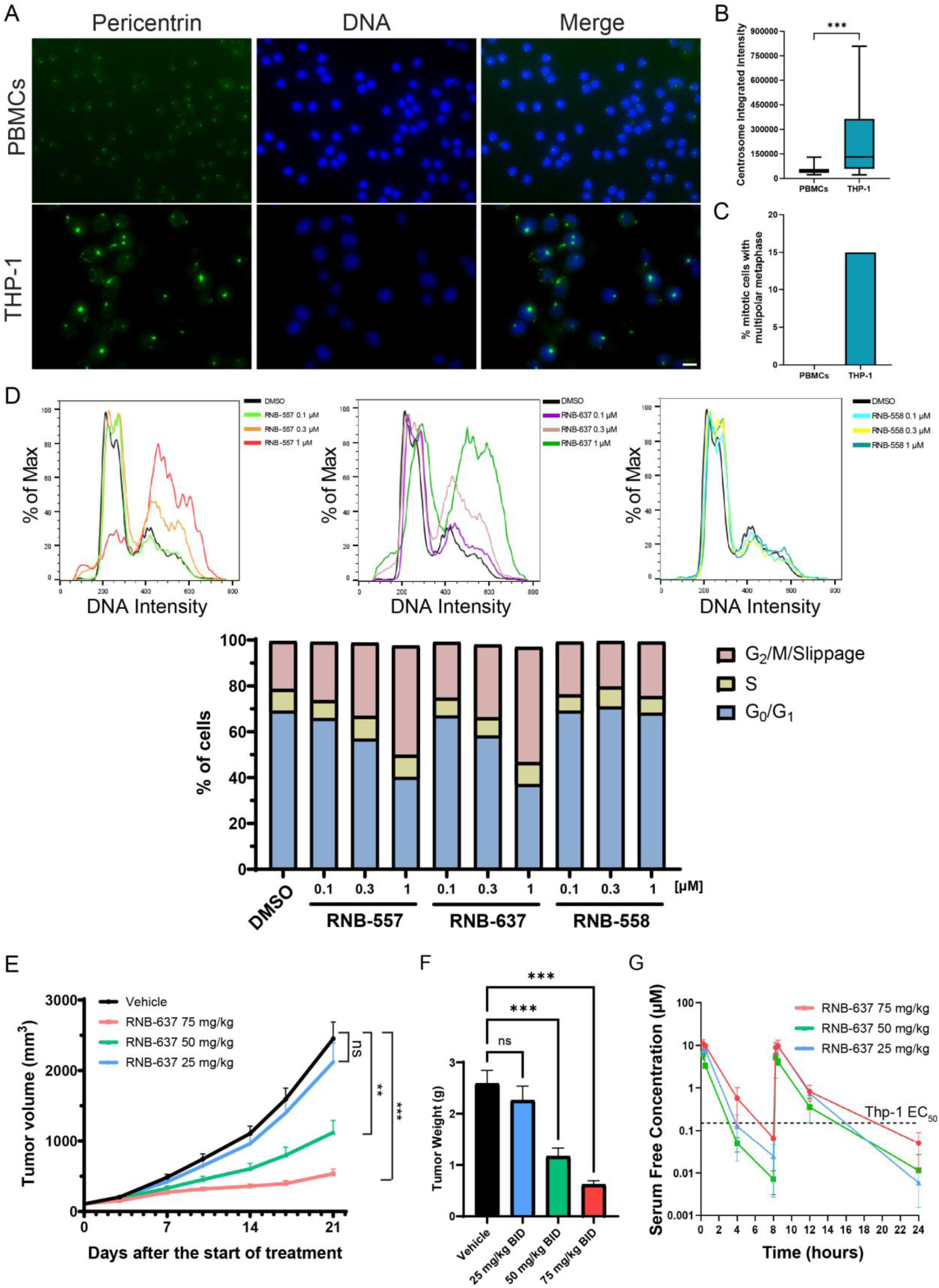
**(A)** Immunofluorescence staining of Pericentrin and Hoechst DNA in PBMCs and THP-1 cells (60×, Z-stack maximum projection). **(B)** Quantification of Pericentrin fluorescence integrated intensity in both cell types. **(C)** Multipolar spindle frequency quantified in THP-1 versus PBMCs (n = 20 mitotic cells per condition). **(D)** Cell-cycle profiling by Hoechst DNA staining and flow cytometry single cell analysis of THP-1 cells following RNB-557, RNB-637 and RNB-558 (inactive isomer negative control) for 14 hours, at the indicated concentrations. Bottom bar graph shows relative cell counts at G0/G1, S and G2/M/Slippage cell cycle phases. **(E)** In vivo THP-1 xenografts in CB17-SCID mice treated with vehicle control and RNB-637 dosed orally, twice per day, at 25, 50, and 75 mg/kg, for 21 days. Mean tumor volume ± SEM is shown (n = 10 mice/group). **(F)** On day 21, tumors were excised, and the mean weight ± SD is shown for the different groups. Two-way Anova p-value is shown (p-value < 0.01 (**), p-value < 0.001 (***), ns-not significant). **(G)** Pharmacokinetic analysis of RNB-637 concentrations following oral administration in CB17-SCID mice. Blood samples were collected at multiple time points over 24 hours from three mice per treatment group on day 21. Mean serum free compound concentration ± SD is shown. The dashed line indicates the Thp-1 EC50 value determined in vitro.

