## Supplementary Information for "FTO regulates centrosome function and mitotic fidelity in cancers with chromosome instability (CIN)"

### Primers

| Target Gene | Forward (5’-> 3’) | Reverse (5’-> 3’) |
|---|---|---|
| FTO | ATTCTATCAGCAGTGGCAGC | AGGTCCCGAAATAAGCAGCC |
| Cyclin B1 | TGAACAACTGCAGGCCAAAA | CCAGCATAGGTACCTTTTCAAGA |
| Cyclin B1 (pre-mRNA) | GCTCTGCCTAACATTTTACATGC | ACCTGTGTGACCCTGACATA |
| HPRT1 | AGGACCTCTCGAAGTGTTGG | TTGCAGATTCAACTTGCGCT |
| GAPDH | TCC ATG CCA TCA CTG CCA CC | TGA CCT TGC CCA CAG CCT TGG |

### siRNAs

Predesigned Mission siRNA targeting FTO (NM_001080432; sequence starting at 1269 nt) was obtained from Merck (Cat# SASI_Hs02_00314786). The non-targeting siRNA was purchased from Dharmacon (ON-TARGETplus Non-targeting Pool, Cat# D-001810-10-05).

### RNA Oligos

RNA oligonucleotides containing repeats of naturally occurring N6-methyladenosine (m6A) RNA motifs[52,53] were custom synthesized by Ella Biotech GmbH. To assess the specificity of m6A-RNA antibodies, the sequences 5′-AUUGUCA(m6A)CAGCAGC-3′ and 5′-AUUGUCACAGCAGC-3′ were synthesized. For the recombinant FTO enzymatic assay and endpoint measurement by ELISA, the following biotinylated oligos were prepared: 5′-Biotin-GG-m6A-CUGG-m6A-CUGG-m6A-CUGG-m6A-CU and 5′-Biotin-GG-A-CUGG-A-CUGG-A-CUGG-A-CU.

**Primary Antibodies**

Primary antibodies used were: anti-FTO (ab126605, Abcam), anti-Cyclin B1 (ab32053, Abcam), anti-Pericentrin (ab28144, Abcam), anti-α-Tubulin (ab7291, Abcam), anti-Aurora B (ab45145, Abcam), anti-NuMA (sc-365532, Santa Cruz Biotechnology), anti-KIFC1 (ab172620, Abcam), anti-Eg5(KIF11) (ab272220, Abcam), anti-TPX2 (ab252944, Abcam), anti-Aurora A (ab108353, Abcam), anti-ALKBH5 [HL2061] (GTX637965, GeneTex), and anti $m^6A$-RNA (ab284130, Abcam).

**Secondary Antibodies**

Secondary antibodies used were: goat polyclonal anti-Rabbit IgG-HRP (ab6741, Abcam), goat polyclonal anti-Rabbit IgG Alexa Fluor 488 (ab150077, Abcam), and donkey polyclonal anti-Mouse IgG Alexa Fluor 568 (A10037, Thermo Fisher).

**Supplementary Information References**

1. Liu, J. et al. A METTL3–METTL14 complex mediates mammalian nuclear RNA *N6*-adenosine methylation. *Nat. Chem. Biol.* **10**, 93–95 (2014).
2. Huang, Y. et al. Small-molecule targeting of oncogenic FTO demethylase in acute myeloid leukemia. *Cancer Cell* **35**, 677–691 (2019).
